# Spatially targeted inhibitory rhythms differentially affect neuronal integration

**DOI:** 10.1101/2024.01.17.576048

**Authors:** Drew B. Headley, Benjamin Latimer, Adin Aberbach, Satish S. Nair

## Abstract

Pyramidal neurons form dense recurrently connected networks with multiple types of inhibitory interneurons. A major differentiator between interneuron subtypes is whether they synapse onto perisomatic or dendritic regions. They can also engender local inhibitory rhythms, beta (12-35 Hz) and gamma (40-80 Hz). The interaction between the rhythmicity of inhibition and its spatial targeting on the neuron may determine how it regulates neuronal integration. Thus, we sought to understand how rhythmic perisomatic and distal dendritic inhibition impacted integration in a layer 5 pyramidal neuron model with realistic dendrites supporting Na^+^, NMDA, and Ca^2+^ spikes. We found that inhibition regulated the coupling between dendritic spikes and action potentials in a location and rhythm-dependent manner. Perisomatic inhibition principally regulated action potential generation, while distal dendritic inhibition regulated the incidence of dendritic spikes and their temporal coupling with action potentials. Perisomatic inhibition was most effective when provided at gamma frequencies, while distal dendritic inhibition functioned best at beta. Moreover, beta modulated responsiveness to distal inputs in a phase-dependent manner, while gamma did so for proximal inputs. These results may provide a functional interpretation for the reported association of soma-targeting parvalbumin positive interneurons with gamma, and dendrite-targeting somatostatin interneurons with beta.

## INTRODUCTION

Cortical circuits are composed of connected networks of excitatory pyramidal neurons. To maintain a balanced level of excitability, they form dense reciprocal connections with local inhibitory interneurons. The two largest subclasses of interneurons in the cortex are parvalbumin positive (PV) and somatostatin positive (SOM) cells [1]. A major distinction between PV and SOM interneurons is that they synapse onto different parts of the pyramidal neuron. PV interneurons tend to contact the soma and proximal dendrites (perisomatic), while SOMs target distal dendrites [2–5]. Numerous studies have identified differences in how they regulate the responsiveness of local excitatory principal neurons [6–8]. These differences could arise from their connectivity, intrinsic properties, or where they synapse onto pyramidal neurons. Indeed, the location of an inhibitory synapse qualitatively changes its effect on synaptic integration [5, 9–11].

Synaptic integration in pyramidal neurons arises from the interplay between passive and active ion channels and dendritic morphology [9, 12–14]. Dendrites produce regenerative spiking events, namely Na^+^, NMDA, and Ca^2+^ spikes. Their initiation depends upon the spatiotemporal coordination of excitatory synapses and interaction with the depolarization of dendritic branches. Incorporating just NMDA spikes into a model neuron radically increases the complexity of its dendritic integration [15]. Na^+^ and NMDA spikes can drive somatic spiking in an *in vivo*-like model neuron [13]. A Ca^2+^ spike in the apical trunk converts a single somatic spike to a burst [9, 16], and may allow pyramidal neurons to act as multi-stage integrators [17, 18]. This complexity is only increased by including inhibition. The emission of bursts of action potentials is controlled by the timing and dendritic location of inhibition [9].

The rhythmicity of inhibition at distal dendrites also affects somatic integration [19]. Reciprocal interactions between excitatory principal neurons and inhibitory interneurons gives rise to rhythmic activities (pyramidal-interneuron network gamma or PING mechanism). To initiate the rhythm, excitatory principal neurons activate inhibitory interneurons that deliver feedback inhibition to principal neurons. This transiently suppresses principal neuron firing. As the inhibition wanes, principal neurons resume their activity and reengage the inhibitory population, starting a new oscillatory cycle. Some evidence suggests that different interneuron subtypes pace rhythms at different frequencies, with gamma oscillations depending on PV [20–22], and beta oscillations on SOM [21]. On the other hand, recent work has also found that beta/low-gamma rhythms in V1 engage both PV and SOM in their generation [23, 24].

Numerous cognitive processes are associated with these inhibitory rhythms in the cortex. Gamma oscillations are especially pronounced during stimulus presentation or behavioral initiation [25–27]. Beta rhythms occur during preparatory states or working memory [28, 29]. But fundamentally, these rhythms probably reflect different modes of local cortical activity and interregional communication [30–32]. For instance, gamma oscillations are associated with feedforward transmission of information in cortical circuits and beta oscillations may mediate feedback [30, 33]; but see [34].

Given all this, it is important to better understand how the various integrative events in the dendritic tree are affected by the location and frequency of inhibition. Perhaps some rhythmic timescales are more effective in the dendrites, and others at the soma. Investigating this is beyond the reach of conventional experimental techniques. Moreover, computational models exploring how beta and gamma influence information processing have mostly relied upon simplified model neurons (e.g., [35, 36]). Thus, using a morphologically and biophysically detailed layer 5 pyramidal neuron model with Na^+^, NMDA, and Ca^2+^ spikes, we investigated how perisomatic and distal dendritic inhibition impacted dendritic integration, and the modulation of these events by beta and gamma rhythmicity.

## RESULTS

### Construction of a model cortical layer 5 pyramidal neuron

To study the effect of inhibitory rhythms on synaptic integration and somatic spiking, we adapted a previously published morphologically and biophysically detailed model of a cortical layer 5 pyramidal neuron (**Fig. 1A**; See methods for details). Modifications to the model were done in accordance with the published literature. In brief, this model featured a multicompartmental dendritic tree that produced dendritic Na^+^, NMDA, and Ca^2+^ spikes, along with somatic action potentials that could backpropagate (**Fig. 1B**). We distributed conductance-based synapses across the dendritic and somatic compartments with an average density of 2.16 excitatory and 0.22 inhibitory contacts per µm. Presynaptic drivers of excitatory synapses were drawn from a pool of 5,200 point process sources that emulated correlated afferent drive with 2-8 synaptic contacts from the same presynaptic neuron. Inhibitory synapses were divided into two populations, those targeting the soma and proximal 100 µm of the dendrites (referred to as perisomatic), and those synapsing outside that area (referred to as distal). To capture excitatory/inhibitory (E/I) balance, a hallmark of cortical activity, the rate of inhibitory synaptic drive was a rescaled version of the rate of excitatory drive, lagged by 4 ms to emulate feedforward inhibition. Naturalistic presynaptic drive elicited a median firing rate of 5.3 Hz, in agreement with *in vivo* rates in cortex (**Fig. 1A** inset, [37]).

**Figure 1:**
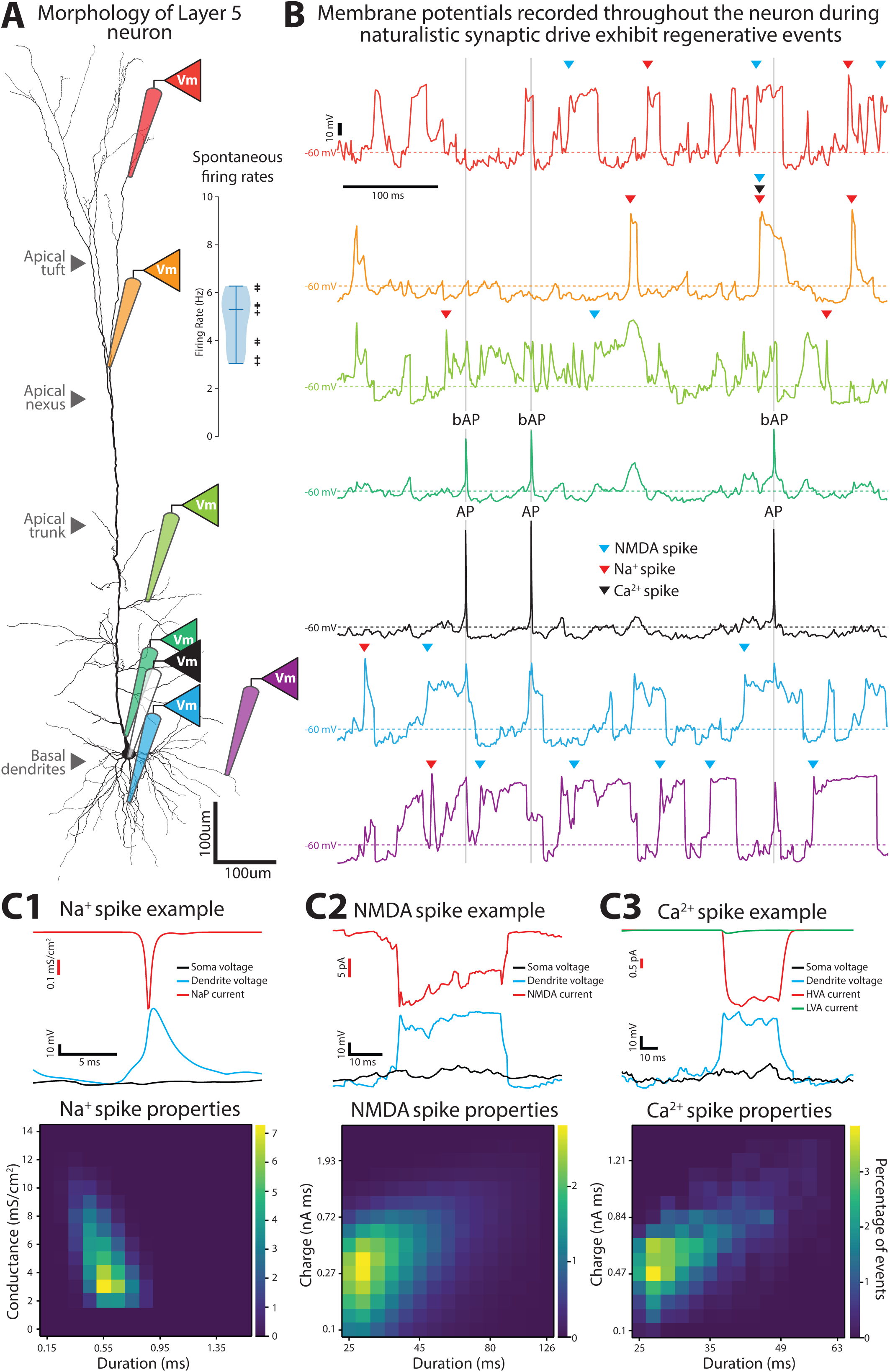
A model layer 5 pyramidal neuron with active dendrites. **(A)** The morphology of the neuron. Virtual recordings can be obtained from any desired compartment (colored pipettes). Inset, naturalistic presynaptic activity drives firing rates in our model like those in vivo. Each black cross is the mean rate for a different simulation. **(B)** Examples of membrane potentials recorded simultaneously across the dendritic tree (in color) and soma (black) during naturalistic drive. Regenerative events are indicated with arrows or text (AP: action potential, bAP: backpropagating action potential). **(C1-3)** Demonstration of our detection of dendritic spike events (top) and characterization of their properties (bottom). Events are binned according to the properties that were used in their detection. Bin edges for the event durations were not evenly set for panels **C2** and **C3**.

Dendrites were endowed with the following voltage-dependent conductances: a fast-inactivating Na^+^ current (I_NaT_), muscarinic K^+^ current (I_m_), fast non-inactivating K^+^ current (I_Kv3.1_), high voltage-activated Ca^2+^ current (I_Ca_HVA_), low voltage-activated Ca^2+^ current (I_Ca_LVA_), and Ca^2+^ activated K^+^ current (I_SK_). As a result, the basal and apical dendrites could generate Na^+^ and NMDA spikes (**Fig. 1B**; [13]). Dendritic Na^+^ spikes were regenerative events lasting less than 1 ms that were not preceded by somatic action potentials (**Fig. 1C1**; [38]). A Na^+^ spike was detected when the dendritic Na^+^ channel conductance (gNa) was higher than 0.3 mS/cm^2^, except when this threshold was reached within 5 ms after a somatic action potential, to distinguish from backpropagating action potentials.

NMDA spikes occur when adjacent NMDA-bearing synapses were synergistically recruited by a combination of glutamatergic activation and local depolarization (**Fig. 1C2**; [18, 39]). They typically lasted between 20 and 80 ms. They were detected when a compartment’s membrane voltage exceeded −40 mV for at least 26 ms and NMDA current exceeded −100 pA ms of charge.

Ca^2+^ spikes are depolarizations generated at the nexus of the apical trunk upon activation of voltage-gated Ca^2+^ channels (**Fig. 1C3**; [40, 41]). They lasted between 20 and 50 ms. To detect them, the membrane potential must exceed −40 mV for at least 26 ms, and the combined Ca^2+^ currents (LVA and HVA) had to be 1.3 times higher than when the voltage criterion was reached (t_v_). The Ca^2+^ spike ended when this value fell to 1.15 times its value at t_v_.

Altogether, under conditions that mirror *in vivo* afferent drive, our model reproduces the dendritic spikes of a layer 5 pyramidal neuron.

### Relationship between dendritic and somatic spikes

Dendritic spikes induced by synaptic activity are the principal drivers of somatic action potentials. Prior experimental and modeling work has found this to be the case for layer 2/3 and layer 5 pyramidal neurons [13, 18, 42–44]. Layer 5 pyramidal neurons have a substantially longer apical trunk, which increases the electrotonic distance of their apical tuft from the soma (**Fig. 2A**), and diminishes the ability of tuft synapses to elicit action potentials. Voltage-gated Ca^2+^ channels at the apical nexus compensate for this by producing a robust Ca^2+^ spike that drives a burst of action potentials at the soma [17, 18]. Underscoring that Ca^2+^ spikes compensate for morphology, pyramidal neurons with shorter apical dendrites have weaker Ca^2+^ spikes that only elicit a single spike [45–47]. We thus assessed how dendritic spikes at different electronic distances from the soma were related to action potential generation.

**Figure 2:**
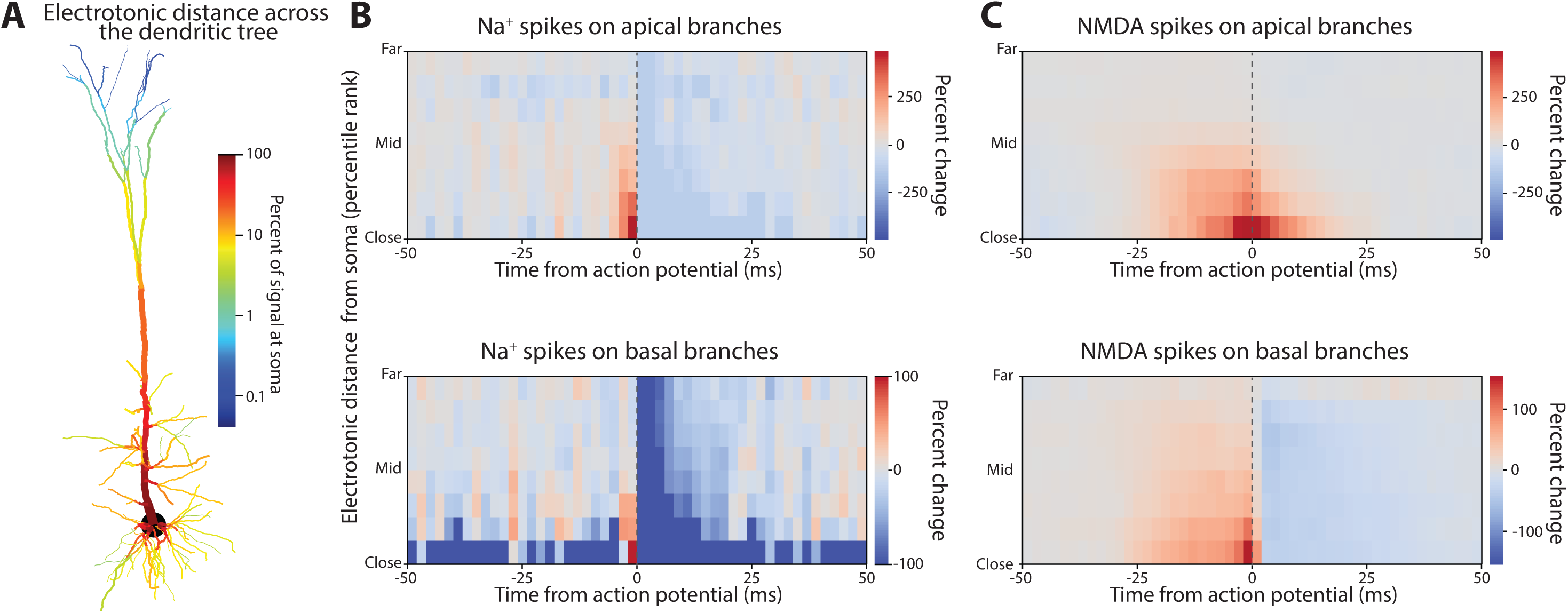
Influence of Na^+^ and NMDA spikes on action potential generation. **(A)** Electrotonic distance between each dendritic compartment and the soma. **(B)** Dendritic compartments were grouped by their type (apical or basal) and electrotonic distance (percentile) from the soma. The percent change in Na^+^ spike presence in those compartments relative to somatic spiking. Na^+^ spikes increased immediately prior to action potentials in dendritic compartments that were electrotonically close to the soma. **(C)** Same format as **B**, but for NMDA spikes. These showed a similar degree of change, but a broader temporal coupling.

Dendritic compartments differed in their degree of passive electrical coupling to the soma (i.e., electrotonic distance; **Fig. 2A**). We measured this by injecting a 20 Hz sinusoidal current in each dendritic compartment, and then calculating the ratio of the membrane voltage response at the soma over that at the dendrite. The apical trunk exhibited a relatively small attenuation ratio of ∼10%. Progressing distally into the apical tuft, attenuation reached 0.1%. Basal dendrites showed greater attenuation than the apical trunk due to their smaller diameter. However, with an attenuation ratio reaching ∼1%, their distal tips were still electrotonically closer than the apical tuft. Thus, large, and long-lasting changes in membrane potential, like those produced by dendritic spikes, would be required to have any effect on the soma.

We examined this by measuring the spike-triggered average between somatic action potentials and the presence of dendritic spikes across the dendritic tree. To simplify the complex geometry of our model neuron, dendritic compartments were grouped into deciles by their electrotonic distance, and whether they were on apical or basal branches. For each time lag from the somatic action potential, we measured the percent change in dendritic spike incidence from the mean rate across the entire simulation.

Dendritic Na^+^ spikes increased 2-3 ms prior to somatic action potentials in both basal and apical dendrites (**Fig. 2B**). This relationship was strongest for the compartments nearest the soma, with the rate of Na^+^ spikes in the apical trunk increasing 300% over baseline prior to somatic action potentials, and 100% in basal compartments. This relationship fell off as the dendritic spikes moved farther away from the soma, indicating that Na^+^ spikes in distal branches had little direct influence on somatic spiking.

The incidence of NMDA spikes increased ∼25 ms prior to somatic action potentials, much earlier than seen with dendritic Na^+^ spikes (**Fig. 2C**). But, like dendritic Na^+^ spikes, NMDA spikes in the apical branch had a stronger coupling with somatic spiking than those in basal branches, and this effect dropped with distance from the soma. NMDA spikes in the apical branch persisted after an action potential, while those in basal dendrites did not, potentially because of the action potential after-hyperpolarization.

Ca^2+^ spikes originate at the nexus, when the apical trunk first branches into the apical tuft. This region is electrotonically close to the entire apical trunk, facilitating the propagation of Ca^2+^ spikes (**Fig. 3A**). In our model, Ca^2+^ spike occurrence increased within 20 ms of somatic action potentials (**Fig. 3B**). Furthermore, we found that NMDA spikes in the apical dendrites tended to precede Ca^2+^ spikes (**Fig. 3C**). Our ionic current-based detection criteria distinguished between these phenomena despite their similar membrane voltage profiles. Since NMDA spikes in the apical tuft normally have a weak relationship to somatic spiking (**Fig. 2C**), they may elicit somatic spiking indirectly by driving Ca^2+^ spikes. Put another way, the apical nexus may serve as a thresholded nonlinearity for NMDA spikes in the apical tuft to drive action potentials [18]. To test this, we measured how a Ca^2+^ spike changed the spike-triggered average between apical tuft NMDA spikes and action potentials (**Fig. 3D, top**). This revealed that action potentials preceded by a Ca^2+^ spike (by up to 20 ms) had increased coupling with apical NMDA spikes. No such change was seen in basal dendrites (**Fig. 3D, bottom**).

**Figure 3:**
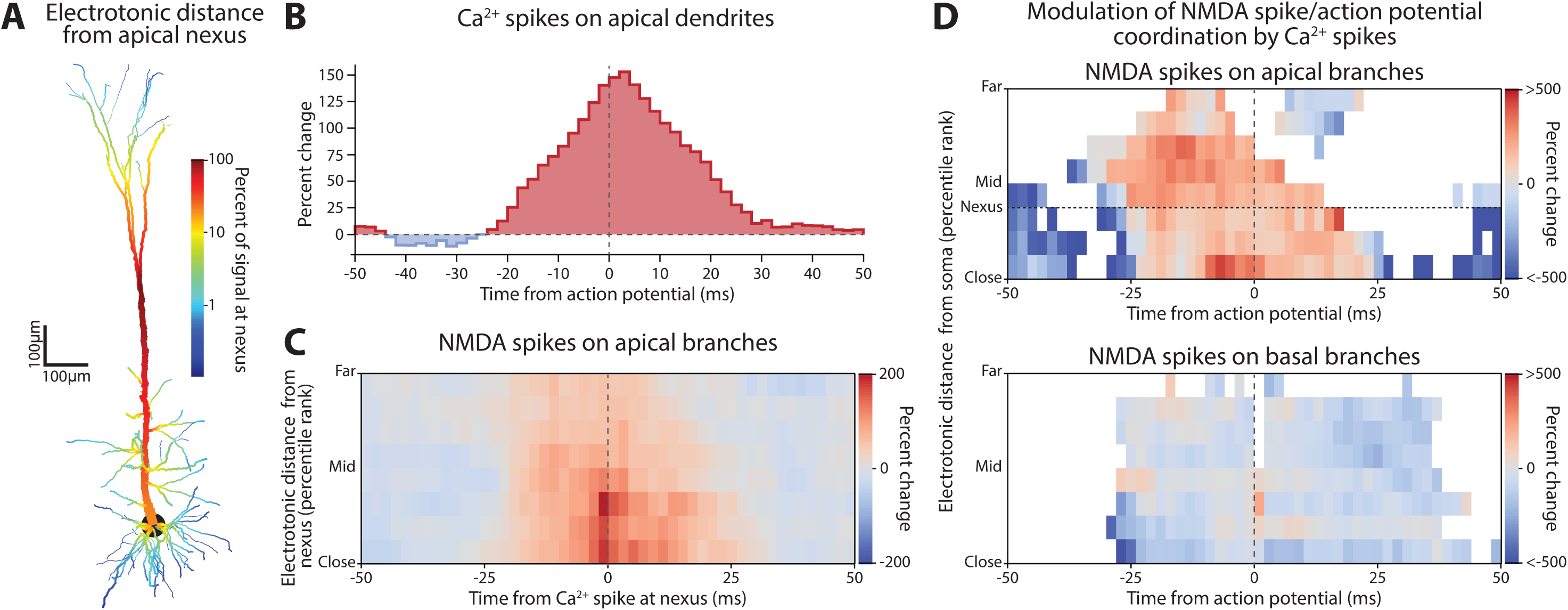
Influence of Ca^2+^ spikes on action potential generation, and their behavior as a second integrative mechanism. **(A)** Electrotonic distance between dendritic compartments and the apical nexus, where Ca^2+^ spikes are generated. **(B)** Change in the incidence of Ca^2+^ spikes at the nexus surrounding action potentials. **(C)** Percent change in NMDA spike presence in the apical dendrites centered on Ca^2+^ spike initiation. **(D)** Percent change in NMDA spike coupling with action potentials during Ca^2+^ spikes. Top, NMDA spikes in apical dendrites were more strongly coupled with action potentials during Ca^2+^ spikes. Bottom, this was not the case for basal dendrites.

### Effect of perisomatic and distal dendritic inhibition on controlling response gain

Subtypes of inhibitory interneurons synapse on distinct dendritic zones. PV interneurons mainly synapse perisomatically, while SOM interneurons target distal dendrites [2–5]. These differences may affect their modulation of synaptic integration [48], either by shifting the threshold for evoking action potentials (a subtractive effect) or altering the slope of the relationship between excitation and firing rate (a divisive effect). There is some disagreement about the degree to which PV and SOM interneurons produce either of these effects [6–8], depending on the source of excitation and local circuitry [49]. So, before we applied beta and gamma rhythmic inhibition in our model, we studied the effect of tonically activating PV- and SOM-like inhibitory synapses.

We doubled the rate of the inhibitory presynaptic drive onto either the PV-targeted perisomatic compartments (somatic and dendritic compartments within 100 µm), or the SOM-targeted distal (>100 µm) dendritic branches. Both decreased the firing rate of the pyramidal cell from 5.5 Hz to less than 1 Hz (**Fig. 4A**; control: 5.5±0.85 Hz; distal: 0.20±0.15 Hz; perisomatic: 0.70±0.31 Hz; mean±SD). These comparable changes may reflect either subtractive or divisive effects and could derive from different mechanisms.

**Figure 4:**
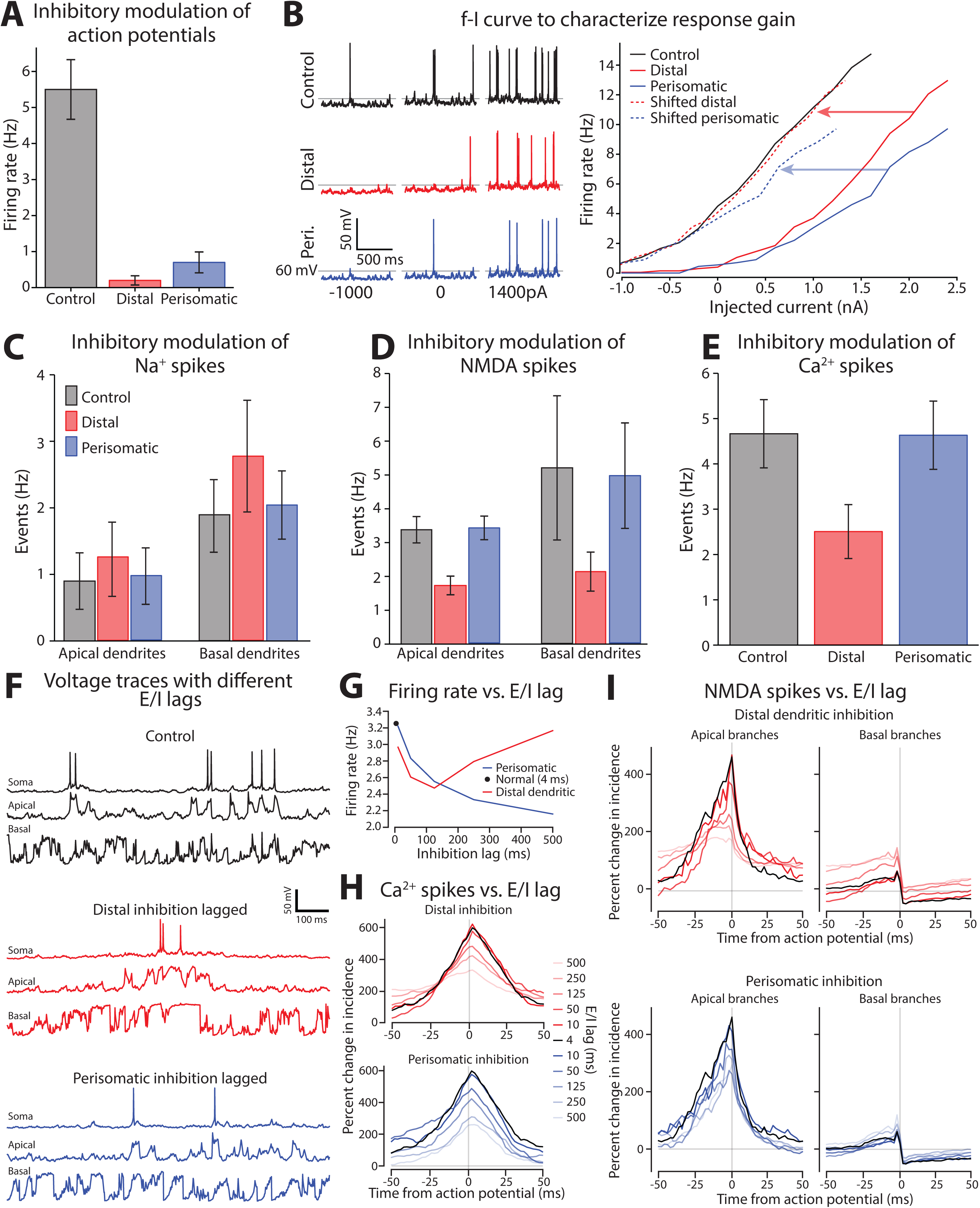
Distal dendritic and perisomatic inhibition reduce action potential generation through different mechanisms. **(A)** Action potential rate during periods with normal inhibitory tone (Control), double rate on Distal branches, or double rate on Perisomatic. Both increases in inhibition dramatically reduced the firing of somatic action potentials. **(B)** Somatic excitability was measured by delivering current steps during the control, distal, and perisomatic inhibition states. Left, example somatic voltage responses to current steps. Right, spike frequency versus current (f-I) curve for each condition. The threshold for evoking an action potential shifted with 2X distal and perisomatic inhibition. Perisomatic, but not dendritic, inhibition changed the f-I slope (compare dashed lines with solid black). **(C)** Impact of altered dendritic inhibition on rate of Na^+^ spikes in apical and basal dendrites. **(D)** Same format as **c**, but for NMDA spikes. **(E)** Rate of Ca^2+^ spikes in the apical dendrites. Basal dendrites lacked Ca^2+^ spikes and were excluded. All error bars are mean ± standard deviation. **(F)** Examples of membrane potential recorded in control (top), and both distal (middle) and proximal (bottom) inhibition lagged by 500 ms. **(G)** Change in firing rate for control (black dot) and for perisomatic (blue) and distal (red) lags in inhibition from 0 to 500 ms. **(H)** Change in incidence of Ca^2+^ spikes for distal (red, top) and proximal (blue, bottom) inhibition. The control case is shown in black in both panels. **(I)** Same as (H) but for NMDA spikes.

To isolate these factors, we first assessed how doubling inhibition affected action potential initiation at the soma. A series of current pulses were injected into the soma to measure the relationship between firing rate and injected current (f-I curve), which captures the gain function of the neuron (**Fig. 4B**). This revealed that both perisomatic and dendritic inhibition shifted the current threshold for action potential initiation. Such an effect is subtractive. In addition, perisomatic inhibition decreased the slope of the f-I relationship compared with the control, which is consistent with a divisive effect.

Although perisomatic inhibition produced the strongest subtractive effect, distal dendritic inhibition reduced firing rate the most (**Fig. 4A**). How can we reconcile these discordant findings? One possibility is that distal inhibition substantially reduces Na^+^, NMDA, or Ca^2+^ spikes. To determine this, we examined the overall rate of dendritic spike events. Perisomatic inhibition did not affect dendritic events compared to the control condition (**Fig. 4C-E**). By contrast, dendritic inhibition decreased NMDA and Ca^2+^ spikes (**Fig. 4D,E**). Na^+^ spikes were relatively unaffected (**Fig. 4C**). This lack of effect may arise from a shortening of the inactivation for voltage-gated Na^+^ channels balancing out the loss of excitatory drive.

### Distinct excitation/inhibition balance effects of perisomatic and distal dendritic inhibition

The previous analysis found distinct effects on neuronal gain and dendritic spiking from tonic changes in perisomatic and distal dendritic inhibition. But *in vivo* inhibition is dynamic and time varying. PV and SOM interneurons form dense interconnections with local pyramidal neurons, supplying time-lagged feedback inhibition. This helps maintain the balance of excitation and inhibition (E/I) in the network by ensuring that an overall increase in excitatory activity is rapidly counterbalanced by proportionate inhibition.

To emulate situations where E/I balance is important, we increased the dynamic variation in excitatory drive (see Methods). As with the previous simulations, the rate of inhibitory synaptic drive was a lagged and rescaled version of the overall excitation rate. Normally that lag is 4 ms, in line with experimental estimates [50]. To probe whether perisomatic or distal dendritic inhibition have distinct effects on E/I balance, we independently varied their lags (**Fig. 4F-I**). One was kept at the nominal 4 ms lag, while the other was extended. Increasing either of their lags by 500 ms produced obvious differences in the emission of dendritic spikes and their coordination with action potentials (**Fig. 4F**). Normally dendritic spikes are evenly distributed in time and drive somatic action potentials. Lagging distal dendritic inhibition clustered dendritic spikes together in time, during which action potentials were emitted. Lagging perisomatic inhibition did not affect the spacing of dendritic spikes, but decreased their coupling with somatic spiking.

We systematically characterized these lag effects for the following spiking events modulated by tonic changes in inhibition: action potentials, Ca^2+^, and NMDA spikes (**Fig. 4G**). Increasing the lag of perisomatic inhibition lowered action potential firing, while for distal dendritic inhibition, the firing rate decreased out to a lag of 125 ms, and then returned to normal at 500 ms. To better understand these effects, we calculated the cross-correlation between dendritic spikes and action potentials. Increasing the lag decreased the coordination between Ca^2+^ and somatic spikes (**Fig. 4H**). For distal dendritic inhibition, this decrease was accompanied by a broadening of their temporal relationship, while a sharp temporal relationship was maintained in the perisomatic case. A similar pattern was observed with NMDA spikes on apical branches (**Fig. 4I**). Basal branches, on the other hand, were relatively unaffected.

In summary, distal and perisomatic inhibition modulate the coupling between synaptic drive and spiking through distinct processes. And while for both cases, the normal 4 ms E/I lag produced maximal conversion of dendritic spiking events into action potentials, extending this lag exerted distinct effects.

### Effect of beta and gamma rhythmic inhibition on neuronal integration

The mechanisms modulating neuronal responsiveness during tonic inhibition of somatic and dendritic compartments may extend to rhythmic inhibition. Thus, we emulated beta and gamma rhythmic input (**Fig. 5A,F**). Depths of modulation were set to similarly entrain action potentials (**Fig. 5B,G**), and were comparable to spontaneous and optogenetically-induced gamma and beta bursts seen *in vivo* [51–54]. Beta rhythmic inhibition was modeled as a 16 Hz sinusoidally modulated rate (20% depth) of the Poisson processes driving inhibitory synapses. Gamma rhythmic inhibition was a 64 Hz sinusoidal modulation (40%) of inhibitory synapses. For both cases, excitatory synaptic drive was a stable Poisson process.

**Figure 5:**
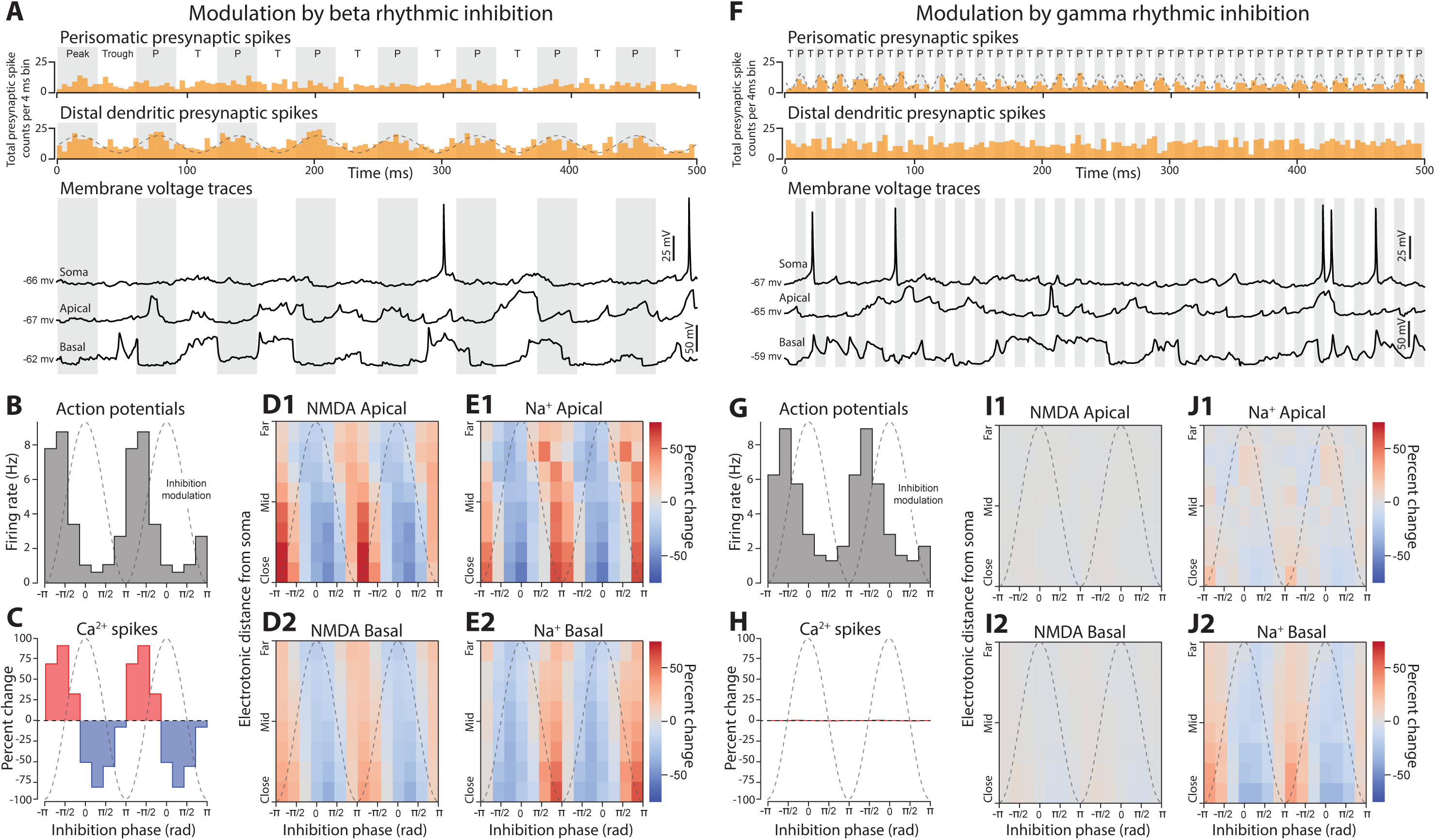
Phase-dependent effects of beta and gamma rhythmic inhibition on dendritic spikes. **(A)** Example data from the beta rhythmic inhibition simulation. Top, presynaptic spike counts. Bottom, voltage traces from somatic and dendritic compartments. Grayed periods are when inhibitory presynaptic spikes are peaking. **(B)** Action potential rate as a function of the phase of the beta rhythm. Dashed gray line shows the modulation of inhibitory drive with respect to phase. **(C)** Percent change in Ca^2+^ spike presence at apical nexus by beta phase. (**D1-2)** Percent change of NMDA spike presence in apical (1) and basal (2) dendrites stratified by electronic distance from the soma. **(E1-2)** Same as **C**, but for Na^+^ spikes. **(F, G, H, I1-2, J1-2)** Same format as above, but with events binned by the phase of gamma rhythmic inhibition. For all graphs phase is given in radians with inhibition at a minimum for −π and maximum at 0.

**Figure 5, Supplementary 1:**
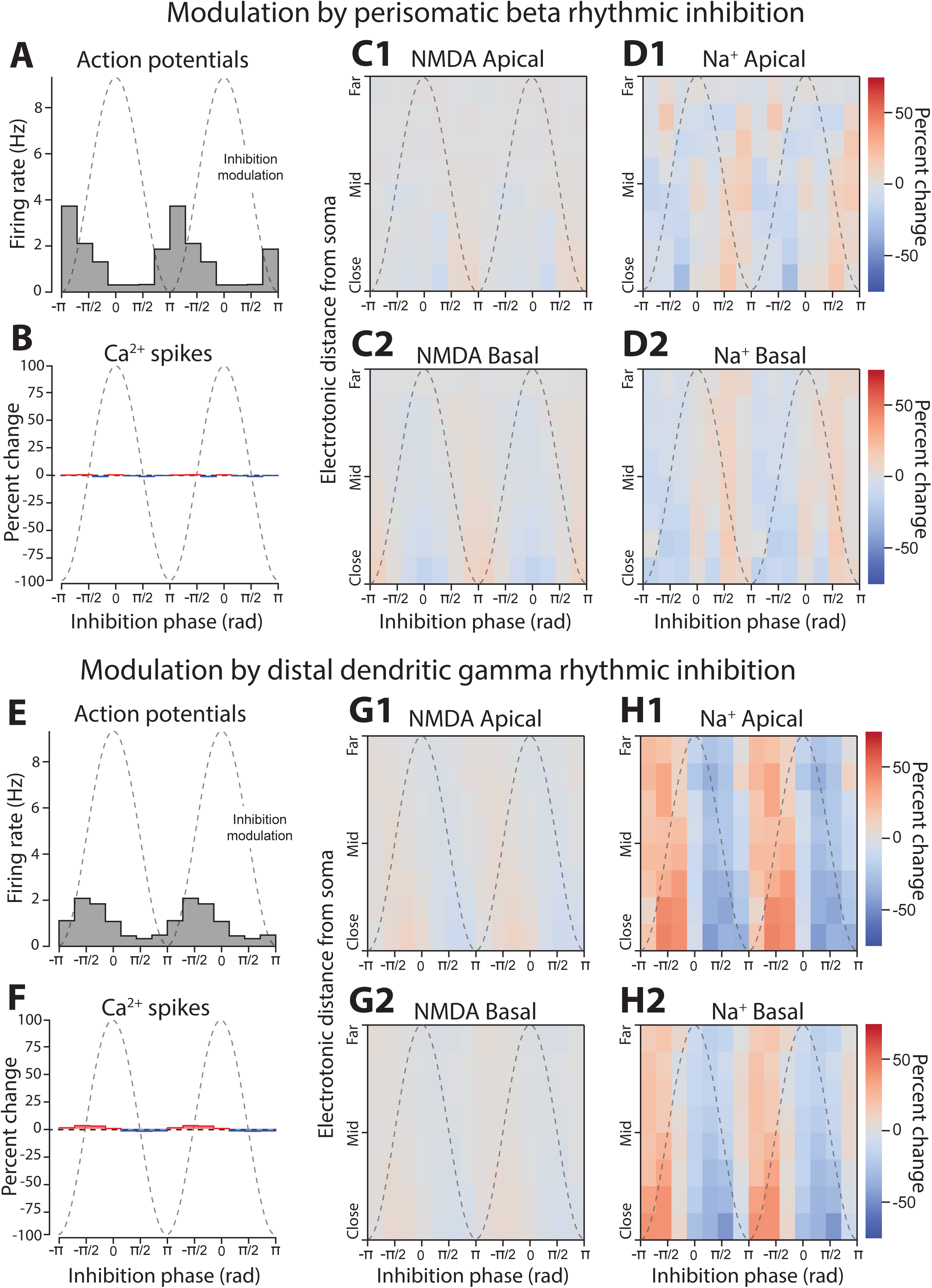
Phase-dependent effects on dendritic spikes of beta and gamma rhythmic inhibition delivered to opposite areas of the neuron. Beta was delivered perisomatically, while gamma was supplied to the distal dendrites. **(A)** Action potential rate as a function of the phase of the beta rhythm. Dashed gray line shows the modulation of inhibitory drive with respect to phase. **(B)** Percent change in Ca^2+^ spike presence at apical nexus by beta phase. (**C1-2)** Percent change of NMDA spike presence in apical (1) and basal (2) dendrites stratified by electronic distance from the soma. **(D1-2)** Same as **C**, but for Na^+^ spikes. **(E, F, G1-2, H1-2)** Same format as above, but with events binned by the phase of gamma rhythmic inhibition. For all graphs phase is given in radians with inhibition at a minimum for -π and maximum at 0.

Initially, we delivered beta rhythmic inhibition to the distal dendritic compartments, and gamma to the perisomatic. Even though both rhythms produced similar depths of modulation of somatic action potentials, the underlying causes were distinct. The phase of beta modulated the occurrence of Ca^2+^, NMDA, and Na^+^ spikes, with each showing an ∼75% depth of modulation with respect to their mean level (**Fig. 5C-E**). These changes were seen across the entire dendritic tree, spanning sites electrotonically close to and far from the soma. In addition, the impact on Na^+^ spikes was unexpected (**Fig. 5E**), since delivery of the same inhibition tonically had little effect. By contrast, gamma had virtually no effect on dendritic spikes (**Fig. 5F-H**). Its strongest impact was on Na^+^ spikes in basal dendritic compartments that were electronically close to the soma.

To uncover the modulation of action potentials by gamma, we turned our attention to somatic action potential initiation (**Fig. 6**). For these analyses we divided the inhibitory rhythm into two phases. During the *peak* phase, inhibition was greater than its mean rate, while during the *trough* phase inhibition was lower (see **Fig. 5A,F**). We found that the somatic action potential voltage threshold shifted lower during the ‘trough’ phase of gamma, when inhibition was at its weakest (**Fig. 6A1**), and without any change in the mean membrane voltage (**Fig. 6A2**). This is consistent with gamma phase modulating the shunting of voltage-gated Na^+^ channel currents, which occurs when GABAergic synapses co-locate with the channels mediating action potential initiation [55]. We also observed changes in action potential initiation to beta rhythmic inhibition, but through a different mechanism. During the ‘peak’ phase of beta, when inhibition was maximal, the threshold for evoking an action potential increased, which may reflect an ‘off-path’ shunting of excitatory current away from the soma and towards the dendrites (**Fig. 6B1;** [56]). Additionally, there was a decrease in membrane voltage during the peak phase, which may correspond to decreased excitation arising from the suppression of dendritic spikes (**Fig. 6B2**).

**Figure 6:**
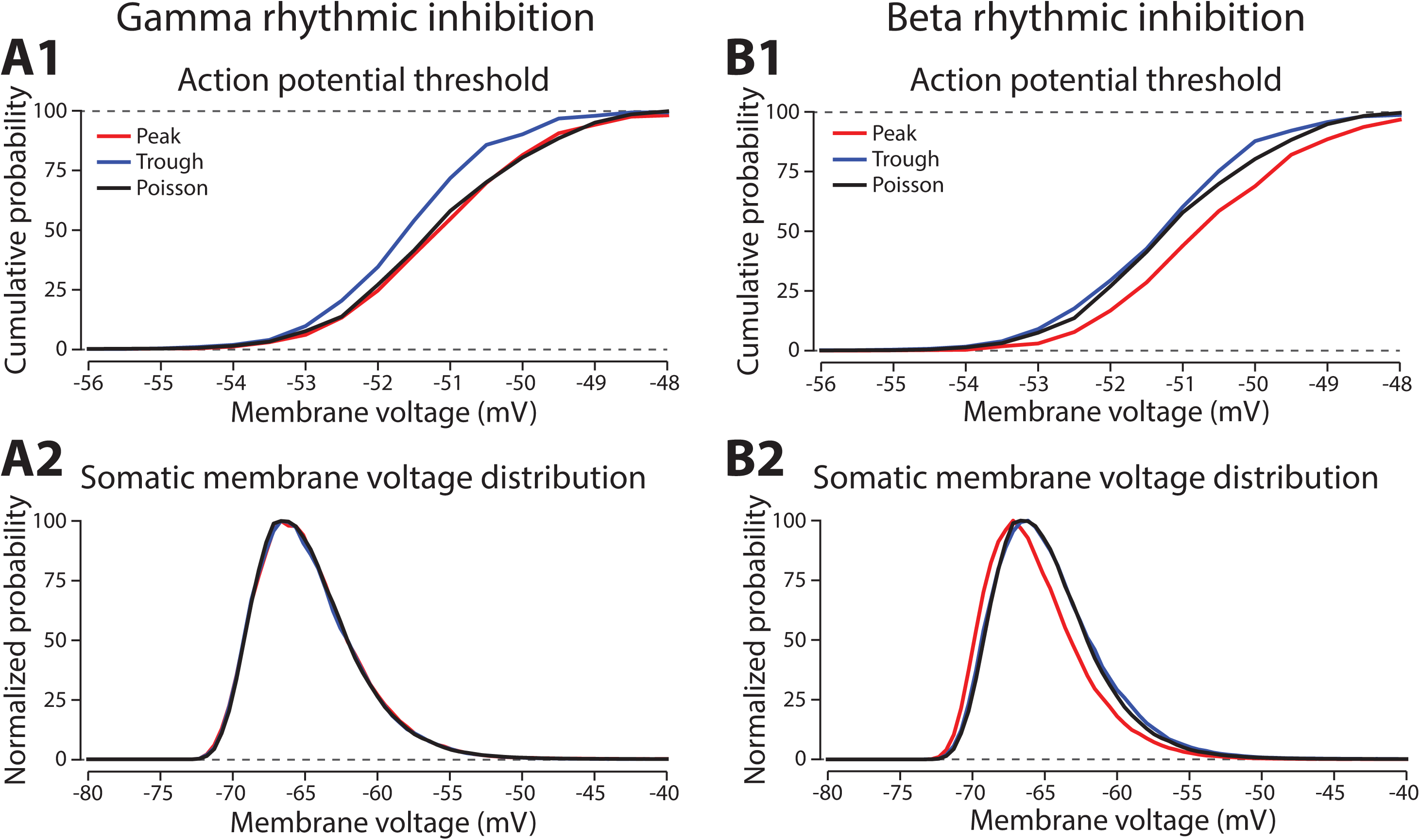
Phase-dependent effects of gamma and beta rhythmic inhibition on somatic excitability. **(A1)** A cumulative probability plot of the distribution of somatic membrane potentials 1ms prior to an action potential, sorted by whether they occurred during the gamma phase with maximal (Peak, red line) or minimal (Trough, blue line) inhibitory drive. Poisson (black) had no rhythmic modulation, but the same mean inhibitory rate. **(A2)** Probability distribution of somatic membrane voltage as a function gamma phase, normalized to the peak probability value. Lines have the same color scheme as in **A1**. **(B1)** Same format as **A1**, but for the beta rhythm. **(B2)** Same format as **A2**, but for the beta rhythm.

**Figure 6, Supplementary 1:**
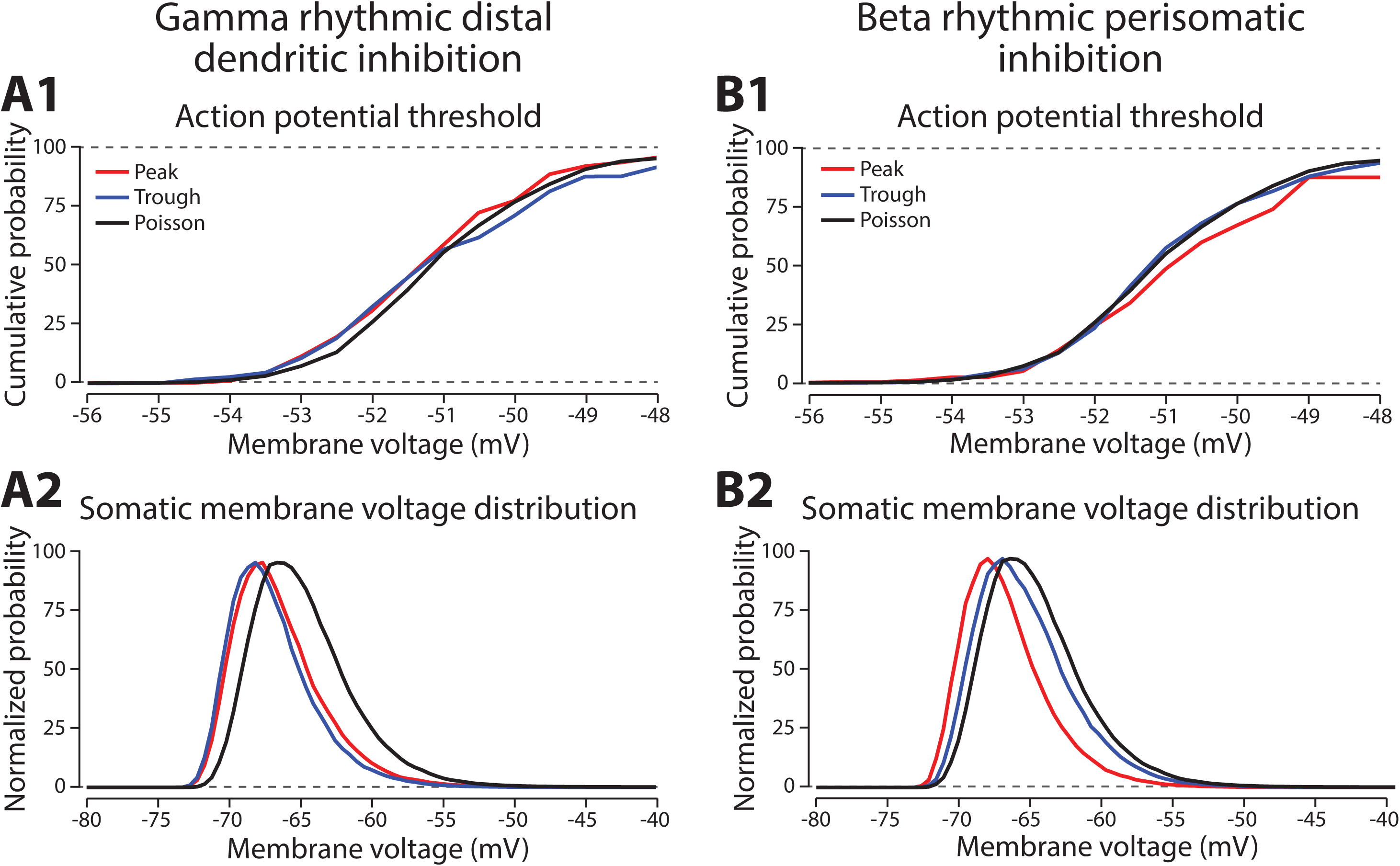
Phase-dependent effects on somatic excitability of beta and gamma rhythmic inhibition delivered to opposite areas of the neuron. Beta was delivered perisomatically, while gamma was supplied to the distal dendrites. **(A1)** A cumulative probability plot of the distribution of somatic membrane potentials 1ms prior to an action potential, sorted by whether they occurred during the gamma phase with maximal (Peak, red line) or minimal (Trough, blue line) inhibitory drive. Poisson (black) had no rhythmic modulation, but the same mean inhibitory rate. **(A2)** Probability distribution of somatic membrane voltage as a function gamma phase, normalized to the peak probability value. Lines have the same color scheme as in **A1**. **(B1)** Same format as **A1**, but for the beta rhythm. **(B2)** Same format as **A2**, but for the beta rhythm.

Accompanying these effects were changes in the rate of dendritic spikes compared with the Poisson inhibition case. Beta increased the rate of Na^+^ (+16.6%) and Ca^2+^ spikes (+15.1%) but decreased the rate of NMDA spikes (−10.7%). Gamma caused no change in NMDA (+0.2%) and a weak increase in Na^+^ spikes (+6.4%), but a robust gain in Ca^2+^ spikes (+37.9%).

We next switched the locations on the pyramidal neuron targeted by the beta and gamma rhythms to disassociate the frequency of inhibition from its location. Beta rhythmic inhibition was delivered perisomatically and gamma rhythmic inhibition to distal dendrites. While phase modulation of firing rate was maintained with both rhythms, the overall level of spiking was dramatically reduced (**Fig. 5S1A,E**). Neither rhythm modulated Ca^2+^ or NMDA spikes (**Fig. 5S1B,C,F,G**). It is likely that the slow time scale of Ca^2+^ and NMDA spikes, ∼50 ms, is not optimal for the fast periodicity of the gamma rhythm, which cycles every ∼15 ms. In agreement with this, Na^+^ spikes, which last less than 1 ms, did show modulation by gamma rhythms delivered to the distal dendrites (**Fig. 5S1D,H**).

Swapping the location of beta and gamma synapses altered their effects on somatic excitability. Gamma rhythmic inhibition on the dendrites had minimal or no impact on action potential threshold, but did shift the somatic membrane potential more negative (**Fig. 6S1A**). This hyperpolarization was not dependent on gamma phase, and likely reflected an overall decrease in the rate of dendritic spikes that could supply excitatory drive to the soma (NMDA: −26.1%, Na^+^: −12.2%, and Ca^2+^: −39.9% compared with the Poisson inhibition case). By contrast, delivering beta rhythmic inhibition to the soma raised the action potential threshold and hyperpolarized the membrane potential during the peak phase (**Fig. 6S1B**). It also reduced the incidence of dendritic spikes (NMDA: −17.1%, Na^+^: −11.5%, and Ca^2+^: −7.6%).

Putting all this together, the effectiveness of rhythmic inhibition depends on where it impinges upon the neuron. Beta rhythms targeting the distal dendrites modulate the incidence of dendritic spikes in a phase-dependent manner, while gamma rhythms delivered perisomatically phase modulate somatic excitability. Swapping the locations of these rhythms diminishes these effects and lowers overall excitability.

### Frequency specific effects of rhythmic inhibition on neuronal integration

Having demonstrated that the location of beta and gamma rhythmic inhibition impacts their effectiveness, we next determined its frequency specificity. To do this, we varied the frequency of rhythmic inhibition between 0.5 and 80 Hz on either the perisomatic or distal dendritic neuronal compartments. Starting with distal dendrites, increasing inhibition frequency above 20 Hz diminished its entrainment of NMDA, Na^+^, and Ca^2+^ spike onsets (**Fig. 7A**). The falloff in entrainment was most pronounced above 20 Hz. Curiously, Na^+^ spikes exhibited a preferential entrainment at 20 Hz. In general, entrainment was strongest in the apical dendrites.

**Figure 7:**
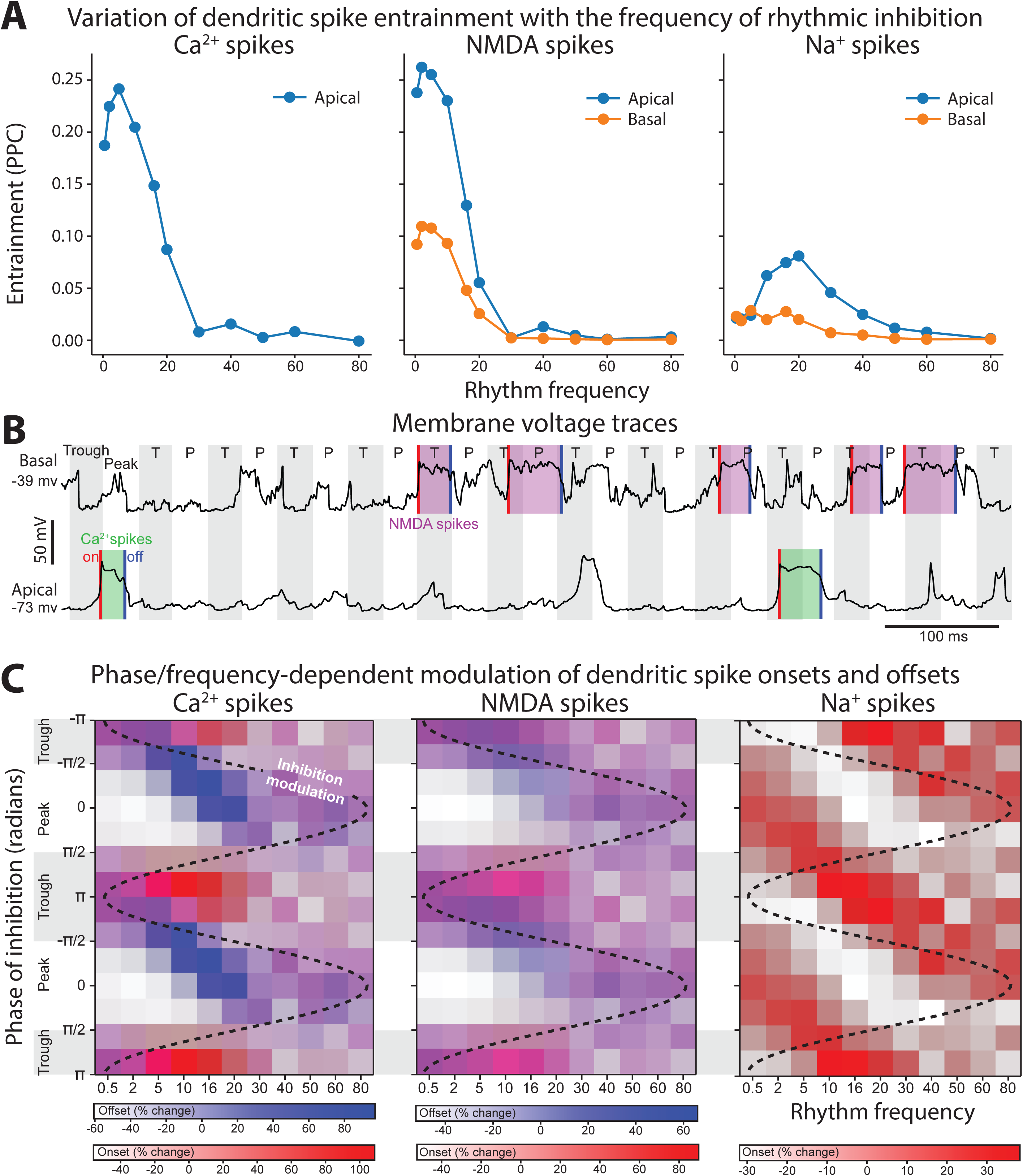
Frequency- and phase-dependent effects of inhibitory rhythms on the distal dendrites. **(A)** Entrainment to an inhibitory rhythm delivered to the distal dendrites varied with its frequency. Higher frequencies were less able to entrain dendritic spikes. Entrainment tended to be strongest for apical (blue) over basal (orange) compartments. **(B)** Example voltage traces from dendritic compartments in either the distal basal or apical branches. Gray shading denotes period where the inhibitory rhythm troughs occurred. For the basal segment, NMDA spikes were shaded in purple, while in the apical segment Ca^2+^ spikes were shaded in green. Dendritic spike onsets denote with red lines, and offsets with blue lines. **(C)** Percent change from the mean in the rate of dendritic spike onsets (red gradient) and offsets (blue gradient) as a function of rhythm frequency and phase. Purple regions denote phase/frequency combinations where both onsets and offsets were elevated, while regions with either just blue or red indicate that offsets or onsets preferentially occurred, respectively. We did not determine an offset for Na+ spikes due to their transience (∼ 1 ms).

The previous analysis considered the entrainment of dendritic spike *onsets*, but NMDA and Ca^2+^ spikes also exhibit *offsets* that could also be modulated by rhythmic inhibition. Indeed, examination of voltage traces in the dendrites during beta rhythmic inhibition revealed that NMDA and Ca^2+^ spike onsets tended to occur during the trough, while offsets happened during the peaks (**Fig. 7B**). To quantify this, we plotted the percent change in the probability of dendritic spike onsets and offsets with respect to both the phase and frequency of the inhibitory rhythm (**Fig. 7C**). For frequencies less than 5 Hz, there was minimal phase separation between the onsets and offsets of Ca^2+^ or NMDA spikes, both events occurred near the rhythm trough, when inhibition was at its weakest. As frequency increased up to 20 Hz, the preferred phase of dendritic spike offsets migrated toward the peak phase, where inhibition is strongest. This phase separation effect was strongest for Ca^2+^ spikes. Thus, beta band frequencies exhibit unique coordination with dendritic spikes: they are the fastest rhythm capable of entraining them and align with their initiation and cessation in a phase-dependent manner.

Turning to perisomatic inhibition, we again varied the frequency of the inhibitory rhythm. Lower frequency inhibition produced phase-dependent shifts in the mean membrane potential (**Fig. 8A**). The trough of inhibition depolarized the soma, while the peak of inhibition had the opposite effect. As the frequency increased, this phase dependent difference went away, vanishing above 50 Hz. By contrast, as frequency increased the bias in momentary changes in the membrane potential diverged between peaks and troughs (**Fig. 8B**). During the peak phase membrane voltage fluctuations were biased negative, while during the trough they were biased positive. This effect increased with frequency, peaking at 50 Hz, and then declining modestly. Together, these effects make gamma frequencies unique in keeping the mean membrane potential equivalent between phases but biasing its fluctuations towards depolarizing or hyperpolarizing with phase. During the trough there was an excess of depolarizing membrane potential fluctuations. Since the rate of excitatory synaptic drive was independent of phase, this suggests that its responsiveness to excitatory inputs increased in a phase-dependent manner with gamma.

**Figure 8:**
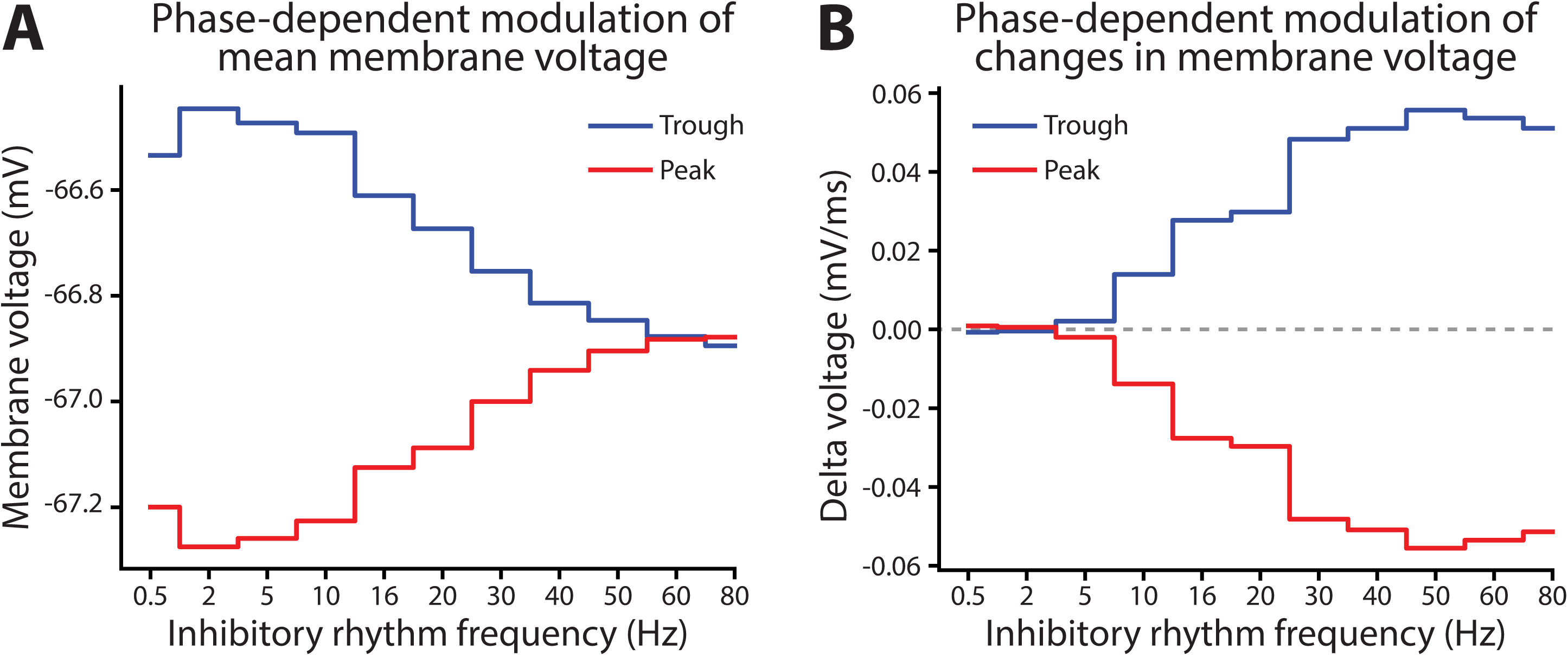
Frequency- and phase-dependent effects of inhibitory rhythms on the perisomatic region. **(A)** The mean somatic membrane potential during either the trough or peak phase of the inhibitory rhythm. **(B)** Mean of the distribution of somatic membrane potential fluctuations as a function rhythm phase and frequency. Fluctuations were measured across the entire simulation time as the difference in membrane potential at 1 ms delays. For both graphs red lines are peaks and blue lines are troughs.

### Modulation of dendritic spikes during oscillatory bursts

*In vivo*, beta and gamma rhythms occur as bursts lasting less than a few hundred milliseconds. Since the previous analyses relied on tonically delivered rhythms, we verified that similar effects were observed with oscillatory bursts. Gamma and beta bursts were delivered to the same model with mean depth of modulation like the tonic case (**Fig. 9A,F**; see Methods for details).

**Figure 9:**
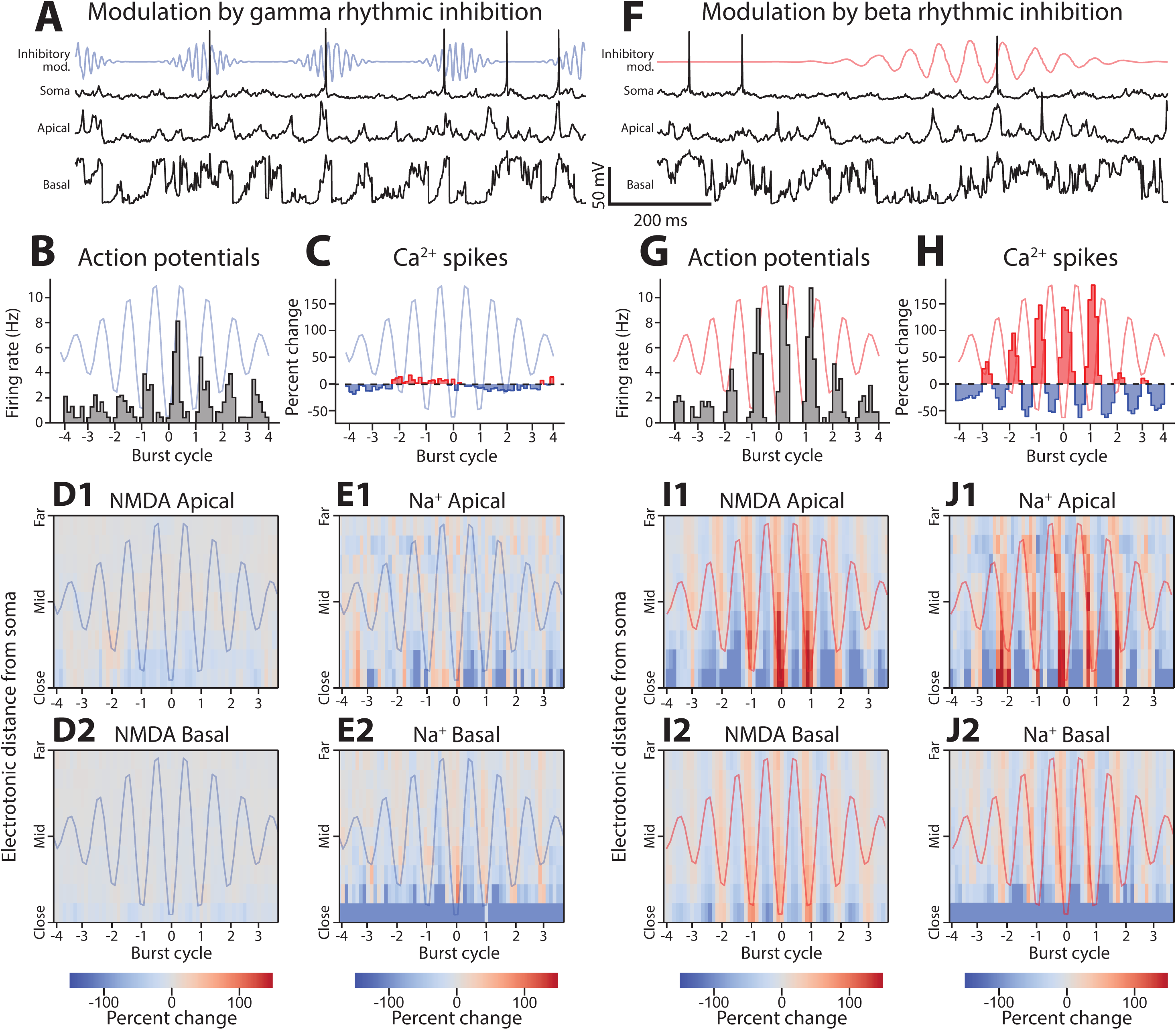
Phase-dependent effects of gamma and beta bursts on dendritic spikes. **(A)** Example data from the gamma rhythmic inhibition simulation. Top, Somatic potential (black line), with firing rate of perisomatic inhibitory synapses (blue line). Middle, voltage trace of apical compartment. Bottom, voltage trace of basal compartment. **(B)** Action potential rate as a function of the phase of the gamma rhythm. Blue line shows the modulation of inhibitory drive with respect to phase. **(C)** Percent change in Ca^2+^ spike presence at apical nexus by gamma phase. (**D1-2)** Percent change of NMDA spike presence in apical (1) and basal (2) dendrites stratified by electronic distance from the soma. **(E1-2)** Same as **C**, but for Na^+^ spikes. **(F, G, H, I1-2, J1-2)** Same format as above, but with events binned by the phase of beta rhythmic inhibition. For all graphs cycle number is given relative to the amplitude peak of the burst.

Under the burst regime, both rhythms mirrored their behavioral effects. Gamma bursts entrained spiking, with entrainment strongest during the middle of the burst (**Fig. 9B**). As with the tonically imposed rhythm, there was none or minimal modulation of Ca^2+^ (**Fig. 9C**), NMDA (**Fig. 9D**), and Na^+^ spikes (**Fig. 9E**). Beta rhythms entrained somatic action potentials (**Fig. 9G**), Ca^2+^ spikes (**Fig. 9H**), NMDA (**Fig. 9I**), and Na^+^ spikes (**Fig. 9J**). These modulations were evident within the first few cycles of a burst, suggesting that they did not require a buildup or evolving entrainment of an underlying process.

### Effect of beta and gamma rhythms on responding to clustered synaptic drive

The results so far suggest that beta and gamma rhythms modulate synaptic integration through different mechanisms depending on their phase and the location of the synapses on the dendritic tree. To examine this further, we added patches of concentrated excitatory synaptic inputs onto either the distal or proximal dendrites (**Fig. 10A**), with densities similar to functional clusters *in vivo* [57, 58]. Coactivated inputs were simulated by driving each synapse with a jittered (2 ms) Poisson process. We ran six separate simulations, either with the distal or proximal clusters engaged, and under conditions of Poisson, beta, or gamma rhythmic inhibition.

**Figure 10:**
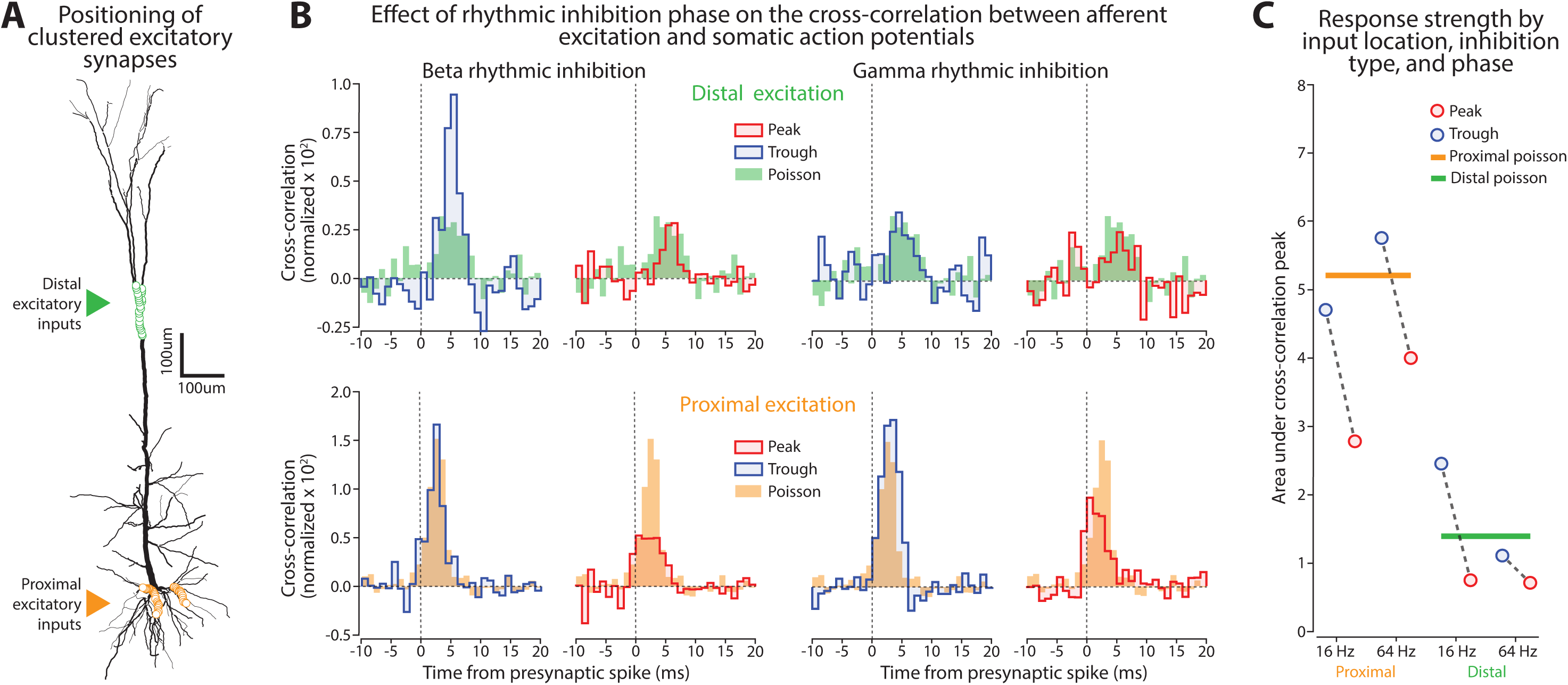
Effect of beta and gamma rhythms on responsiveness to synaptic inputs targeting distinct regions of the dendritic tree. **(A)** Schematic of the location for clustered excitatory synaptic inputs. **(B)** Normalized cross-correlation between synaptic drive onto a clustered input and spiking at the soma, stratified by whether the presynaptic spike arrived during the Peak (red line) or Trough (blue line) of the rhythm. Solid bars correspond to the Poisson stimulation case where inhibition was not rhythmically modulated. Top left, effect of beta on distal inputs. Top right, effect of gamma on distal inputs. Bottom left, effect of beta on proximal inputs. Bottom right, effect of gamma on proximal inputs. **(C)** Summary of effects in panel **B** where the strength of each normalized cross-correlation was measured as its area under the curve. Dots are connected by dashed gray lines if the data points came from the same simulation but at different phases of the rhythm. Solid horizontal lines reflect the cross-correlation strength in the Poisson inhibitory case (no rhythmicity).

To capture how beta and gamma influence synaptic integration we measured the cross-correlation between presynaptic activations at the clustered input and somatic spiking. Separate cross-correlograms were calculated depending on whether the presynaptic spikes occurred during the peak or trough phase of the inhibitory rhythm. This necessarily introduced spurious periodicities into the cross-correlogram that were compensated (see Methods for details). Relative to the arhythmic Poisson inhibition case, beta rhythms enhanced the transmission of distal inputs when inhibition was low (trough phase) and suppressed them when inhibition was high (peak phase, **Fig. 10B, top left**). Proximal inputs were either unaffected or moderately suppressed during the trough and suppressed during the peak (**Fig. 10B, bottom left**). The opposite was the case for gamma. It barely affected or moderately suppressed distal inputs (**Fig. 10B top right**), while proximal inputs were enhanced during the trough and suppressed during the peak (**Fig. 10B bottom right**).

Summarizing these results (**Fig. 10C**), we found that somatic spiking driven by clustered proximal synapses was bidirectionally modulated by gamma rhythms and suppressed by beta. On the other hand, spiking driven by distal clusters was bidirectionally modulated by the beta rhythm and suppressed by gamma. Thus, both rhythms regulate the sensitivity of pyramidal neurons to afferents throughout the dendritic tree, but in a counterposed location-dependent manner.

## DISCUSSION

Arising from multiple interneuron subtypes, inhibition sculpts pyramidal neuron activity by acting at different membrane regions and distinct rhythmic frequencies (**Fig. 11A**). Little was known about how these factors interact with the complexity of dendritic integration. To address this, we characterized the interaction between the location and rhythmicity of inhibition on integration in a morphologically and biophysically detailed layer 5 pyramidal neuron model with active dendrites. We found that distal dendritic inhibition modulated the occurrence of dendritic spikes, while perisomatic inhibition altered action potential generation. This translated into location-specific differences in the effectiveness of inhibitory rhythms. Beta rhythmic inhibition entrained dendritic spikes, focusing them into the phase when inhibition was at a minimum, but only when delivered to the distal dendrites (**Fig. 11B**). In contrast, gamma modulated the threshold for action potential initiation, but only when delivered perisomatically (**Fig. 11C**). The effects of these rhythms were frequency specific, with the timing of beta and gamma aligning preferentially with phase dependent effects on neuronal integration. As a likely result, beta oscillations bidirectionally controlled transmission in distal dendrites and suppressed those onto proximal dendrites, while gamma oscillations did the opposite.

**Figure 11:**
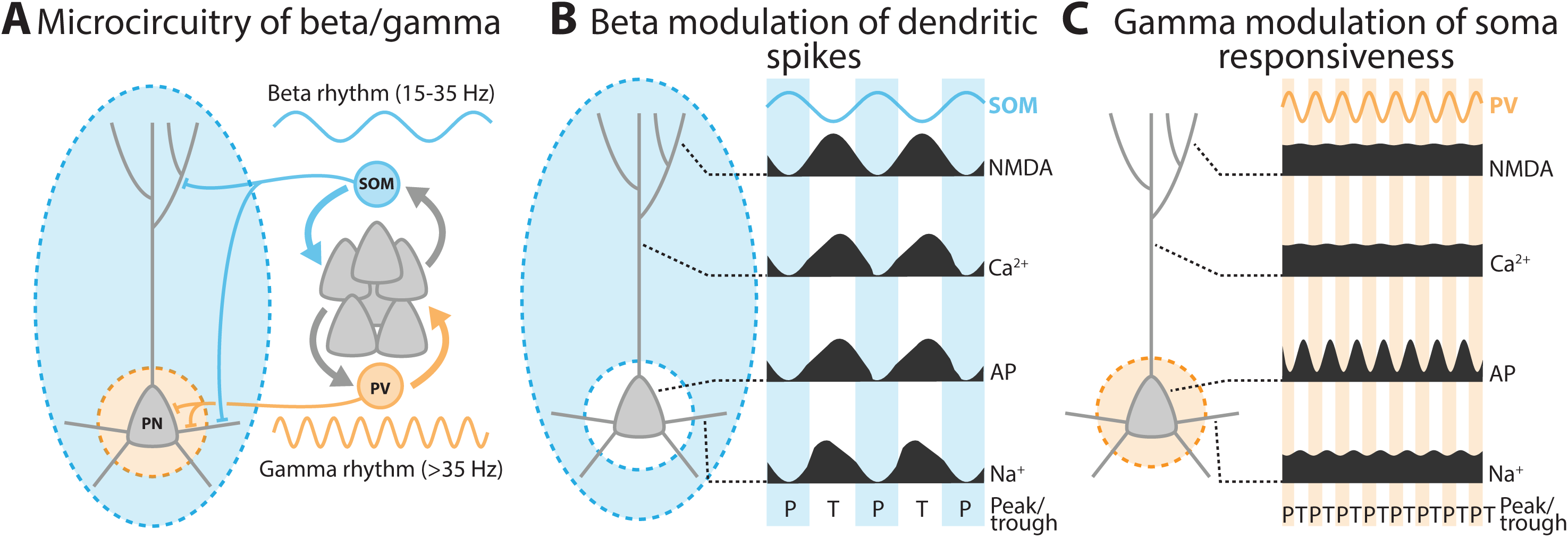
A summary schematic of the principal findings. **(A)** The microcircuitry that was simulated in this study. **(B)** Beta rhythmic inhibition to the distal dendrites modulated dendritic spikes. **(C)** Gamma rhythmic inhibition to the perisomatic region modulated action potential initiation. AP stands for action potentials.

This indicates that the spatial targeting and timing of inhibition go hand in hand. Their alignment is thus fundamental to their distinct effects on neuronal integration. And in turn, this affects function at multiple levels, from synaptic responsiveness to neural coding, and microcircuit operation.

### Neuronal responsiveness

Oscillatory rhythms potentially affect synaptic integration through several mechanisms. Early theorizing held that oscillations synchronize populations of excitatory neurons such that their efferent synapses benefit from spatiotemporal summation in eliciting action potentials [59]. Later it was proposed that if neurons in two regions exhibited coherent variations in their somatic membrane potentials, bringing them closer to or farther from action potential threshold, they could preferentially exchange activities [60]. Modeling and *in vitro* studies have shown that the phase of a sinusoidal current injected at the soma modulates the timing of evoked action potentials [61] or their probability of occurrence [62]. Similar gamma phase dependence has been found *in vivo* [63].

Rhythmic inhibition may also periodically modulate the sensitivity of the neuron to synaptic input. Inhibitory synapses achieve this by lowering the membrane resistance i.e., shunting [64]. This makes it harder for an excitatory synaptic current to drive the membrane voltage toward the action potential threshold. Indeed, optogenetically stimulating PV interneurons at gamma frequencies modulates the responsiveness of cortical neurons to sensory inputs in a phase-dependent manner [65]. However, normally PV interneurons are driven by local principal cells, so the inhibition they impose reflects the aggregate activity in the local network (leading to E/I balance, [66]). Model networks wired in this manner emit spontaneous gamma bursts that produce phase-dependent modulation of the relationship between EPSP amplitude and spiking probability, with a strong positive relationship during the trough of gamma, and a weak one during the peak [36]. Consistent with these results, putative monosynaptically connected pairs of single units recorded *in vivo* show an increase in spike transmission probability during the trough of gamma [67].

To summarize the present possibilities, oscillations could modulate a neuron’s responsiveness by (1) synchronizing the synaptic inputs impinging upon it, (2) modulate how close the membrane potential is to the action potential threshold, or (3) vary its sensitivity to synaptic inputs via shunting inhibition.

Our results fill out this picture in two ways. First, we add a fourth mechanism, which is (4) the modulation of dendritic spiking events. Second, we found that beta and gamma oscillations differentially engaged these mechanisms. Beta oscillations primarily caused 2 via 4, and affected 3. Gamma, on the other hand, operated primarily through 3. It should be noted that to isolate these effects, our model had Poisson excitatory synaptic inputs, which precluded mechanism 1, afferent synchrony. However, it is likely the degree of synchrony would impact both rhythms. Supporting this, *in vivo* whole-cell recordings from neurons in mouse visual cortex exhibit brief membrane depolarizations in synchrony with gamma [68].

### Relevance to coding

Given that beta and gamma rhythms influence spiking via distinct mechanisms, they may also differentially impact coding. During gamma bursts in V1, neurons show a stronger modulation of their firing rate by the contrast of grating stimuli [69]. Gamma phase also modulates orientation selectivity and correlated variability between neurons [70]. This enhancement is modulated by attention [71]. Our results indicate that these effects reflect shifts in somatic sensitivity to excitatory synaptic inputs.

Much less is known about the influence of beta oscillations on coding. In the visual cortex of mice, activation of SOM interneurons increased with the visual stimulus size and homogeneity ([22], note that this study referred to the rhythm as low gamma). In a different study, using a sequence working memory task in primates, the order of presented items corresponded to their phase in the beta rhythm [72]. However, this does not indicate how coding varies with beta phase. Considering our finding that beta synchronized dendritic spikes, it may operate by enabling the summation of normally asynchronous afferents on distal dendrites, facilitating their ability to drive somatic spiking.

One interpretation of rhythms arising from local inhibitory feedback is that they maintain the balance between excitation and inhibition. This can be thought of as a normalization operation that maintains activity within a set range. Normalization can be achieved either through a subtractive effect that raises the threshold for initiating an action potential, or a multiplicative effect that lowers the slope of the relationship between excitation and action potential firing rate. When considered at the population level, these normalization effects impact coding in different ways. Subtractive normalization increases sparsity by dropping out neurons whose excitation is below the raised threshold. Multiplicative normalization, however, encourages dense codes by scaling down firing rates and compressing the range of firing rates. This study found that while both perisomatic and distal dendritic inhibition produced subtractive effects, only perisomatic had a multiplicative effect. Tying this to beta and gamma, beta rhythms may encourage sparse population codes while gamma allows for dense.

### Interaction with microcircuitry

Pyramidal neurons are embedded in a cortical column where they are sparsely connected amongst themselves and densely connected with local interneurons. The dense connectivity with interneurons is crucial for the generation of local beta and gamma rhythms [21, 22]. How might this circuitry interact with the differential regulation of dendritic integration by beta and gamma?

We found that beta rhythms preferentially modulated a pyramidal neuron’s response to inputs on its distal dendrites. The apical tuft in the superficial layers of cortex primarily receives long-range inputs from regions higher up in the cortical hierarchy [73, 74]. This makes beta rhythms ideally positioned to modulate these top-down/feedback signals. Since the rhythm likely derives from local activation of SOM interneurons, it is worth considering what situations would activate those and how they might relate to the functioning of apical dendrites. SOM interneurons receive facilitating synapses from local pyramidal neurons [75, 76], which would make them especially sensitive to bursts of action potentials arising from Ca^2+^ spikes at the apical nexus [18]. Since Ca^2+^ spikes are driven by synaptic activation in the apical tuft, it is likely that beta rhythms regulate the generation of action potential bursts arising from long-range inputs.

Particularly in the visual cortex, SOM interneurons can generate a rhythm in the 25-30 Hz range [22]. We found this to be at the upper end of the frequency range for dendritic inhibitory rhythms to be effective in modulating NMDA and Ca^2+^ spikes. If this rhythm solely recruited SOM interneurons, its effectiveness would be marginal. Potentially compensating for this, recent work has found that PV interneurons also participate in beta/low-gamma [23, 24] (but see [21, 22]). In our model, on its own when beta rhythmic inhibition was delivered perisomatically we found that it was less able to entrain spiking and had an overall hyperpolarizing effect. However, if delivered in conjunction with the distal dendritic inhibition arising from SOM interneurons, this may strengthen entrainment.

Turning to gamma, our results indicated that it mainly affects processing in the soma and proximal dendrites. These are preferentially targeted by layer 5 pyramidal neurons, which sparsely interconnect with projection-type specificity [77–79]. A confluence of evidence implicates PV interneurons in the production of high-frequency inhibitory rhythms such as gamma [80, 81]. Given that PV interneurons diffusely interconnect with local pyramidal neurons [82, 83], and the synapses they receive from them are depressing [75, 76], they likely regulate abrupt increases in local ensemble activity. Interactions within ensembles that precede activating the PV population will be boosted, while those following will be attenuated.

### Interneuron specializations and rhythm timescales

Lastly, it is worth considering why beta and gamma rhythms are primarily mediated by different types of interneurons (but see [24]). One possibility is that the differing pace merely arises from the electrotonic timescales of their feedback inhibition; SOM cells target distal dendrites and thus produce a slower inhibitory feedback signal, versus PV cells delivering rapid perisomatic inhibition. Indeed, we found that swapping the cellular compartments receiving beta and gamma rhythmic inhibition impaired their effectiveness and lowered overall excitability.

So, while our results suggest that spatial targeting of SOM and PV interneurons aligns with the timescales of their *network-level* rhythms, it could also be that their timing and subcellular localization interact to produce specialized *neuron-level* functions [84]. For instance, NMDA and Ca^2+^ spikes in the distal dendrites last for ∼50 ms, making the slower beta rhythm more appropriate for bidirectionally controlling them. Both can be described as dynamical systems with distinct phases with differing sensitivity to inhibition. Ca^2+^ spikes are dynamical events comprised of an initiation, plateau, and termination phase. Inhibition delivered during the plateau phase shortens their duration [85]. If the beta rhythm is comprised of cycling between periods of elevated excitation (increased NMDA spike generation) followed by elevated inhibition, then Ca^2+^ spike initiation will tend to occur during the excitatory phase, and its plateau during the subsequent inhibitory phase. A plateau during the inhibitory phase will more quickly enter termination. This is bidirectional control. On the other hand, slower rhythms (e.g., 1 Hz) initiate Ca^2+^ spikes during the excitatory phase that plateau and enter termination autonomously, before the inhibitory phase is reached. The same principle holds for NMDA spikes [86]. As a result, rhythms in the range from 15-30 Hz are optimal for synchronizing the onsets and offsets of dendritic spikes across a population of neurons.

The integrative effects of gamma (>40 Hz) are also specialized. Low frequency inhibitory rhythms delivered to the soma tended to shift the membrane potential higher or lower with the rhythm’s phase, effectively bringing it closer or farther from action potential threshold but not changing the neuron’s sensitivity to fast synaptic inputs. In the gamma frequency range, this is reversed, with the mean membrane potential not varying with rhythm phase but with a shifting bias to positive or negative membrane potential fluctuations. In addition, the trough phase of gamma lowers the threshold for action potential initiation, while slower rhythms like beta only raise the threshold. Consequently, the timing of gamma is ideal for increasing the sensitivity of the neuron to rapid excitation. This agrees with the observation that gamma oscillations accompany rapid excitation-inhibition balancing [87].

## Conclusion

For the most part, the study of beta and gamma rhythms has focused on their correlation with task-related events and entrainment of neuronal firing. Fundamentally, these effects derive from their influence on neuronal integration, which has multiple stages, from modulating the post-synaptic response, inducing a dendritic spike, and triggering an action potential. Since beta and gamma rhythms differentially impacted these processes, this invites a reappraisal of their role in coding and communication, along with unveiling a new hypothesis space for understanding their function.

## METHODS

We adapted a previously published model of a L5 pyramidal neuron [12, 88] to the Brain Modeling Toolkit [89] format to facilitate reproducibility. This open-source software utilized NEURON 7.7 [90] with an integration time step of 0.1 ms for simulating the membrane potential. We added synapses and designed inputs in accordance with the literature to study synaptic integration in vivo. Details of the passive membrane properties and active conductances are only described briefly (full details can be found in [88]). Our focus here is on describing the enhancements we incorporated to mimic in vivo features in the model neuron.

### L5 pyramidal cell compartmental model

Briefly, the cell was built on a morphological reconstruction of a L5 pyramidal tract neuron and included 10 active Hodgkin-Huxley conductances with kinetics taken from the literature [2].

These channels were a fast-inactivating Na^+^ current (I_NaT_), persistent Na^+^ current (I_NaP_), non-specific cation current (I_h_), muscarinic K^+^ current (I_m_), slow inactivating K^+^ current (I_Kp_), fast inactivating K^+^ current (I_Kt_), fast non-inactivating K^+^ current (I_Kv3.1_), high voltage-activated Ca^2+^ current (I_Ca_HVA_), low voltage-activated Ca^2+^ current (I_Ca_LVA_), and Ca^2+^ activated K^+^ current (I_SK_). The somatic compartments contained all of these channels except I_m_. Both the apical and basal dendrite compartments contained I_Ca_LVA,_ I_Ca_HVA,_ I_SK,_ I_Kv3.1,_ I_NaT,_ I_m,_ and I_h_. The membrane capacitance was set to 1 μF/cm^2^ for the soma and axon and 2 μF/cm^2^ for the basal and apical dendrites to correct for dendritic spine area. Axial resistance was set to 100 W-cm and leak reversal potential was −90 mV.

### Divergence, probability of transmission, and functional clustering

Experimental studies indicate that each presynaptic excitatory neuron forms 2-8 synapses with its pyramidal partners, with a release probability between 0.16 and 0.9 [91]. Further, the 2-8 synaptic contacts occur over a span of 20-100 μm [92]). We constrained our model to have these properties. In addition, since synapses may cluster on dendritic branches such that functional presynaptic cell assemblies activate the same branch within 5-10 μm [93, 94], we forced synapses coming from the same presynaptic cell assembly to cluster in this way. In summary, a single presynaptic input fiber could have 2-8 synapses, drawn from a uniform distribution with a release probability between 0.16 and 0.9, also drawn from a uniform distribution 91]. And the synapses from a functional group formed clusters within 5-10μm on the same dendritic branch over a span of 100 μm.

### Synaptic conductances

We randomly distributed excitatory synaptic conductances following previous modeling work [13]. Maximum conductance values followed a log-normal distribution with a mean of 0.2 nS and standard deviation of 0.345 nS for excitatory synaptic conductances. Inhibitory synaptic conductances were fixed at 1 nS [12]. We validated these conductances against excitatory and inhibitory post-synaptic currents measured *in vitro.* The model’s values are compared with experimental values in Table 1.

**Table 1.** Inputs to L5 PN. L5 PN dendrites can course up to layer 1 and receive both excitatory and inhibitory inputs to their dendrites. Here we quantified, where possible, the experimental values for synaptic magnitude, firing rate, divergence, and release probability. We matched the model parameters to the experimental values as closely as possible while preserving a reasonable basal firing rate.

| Synapse type | Characteristics | Model | Experimental |
| --- | --- | --- | --- |
| Excitatory (Basal) | Magnitude of EPSCs | $37.0 \pm 32.3$ pA | **$30.6 \pm 29.9$ pA** [78] |
| | Firing rate | $4.43 \pm 2.9$ Hz | **$4.43 \pm 2.9$ Hz** (Headley personal comm.) |
|  | Divergence | 2-8 | **2-8** [92, 100, 107] |
|  | Number of synapses | 10042 | **10042** [95] |
| | Release probability | $0.53 \pm 0.22$ | **$0.53 \pm 0.22$** [108] |
| Excitatory (Apical) | Magnitude | $25.9 \pm 24.9$ pA | **$30.6 \pm 29.9$ pA** [78] |
| | Firing rate | $4.43 \pm 2.9$ Hz | **$4.43 \pm 2.9$ Hz** (Headley personal comm.) |
|  | Divergence | 2-8 | **2-8** [92, 100, 107] |
|  | Number of synapses | 16070 | **16070** [95] |
| | Release probability | $0.53 \pm 0.22$ | **$0.53 \pm 0.22$** [108] |
| Inhibitory (Perisomatic and somatic) | Magnitude | $162.5 \pm 103.1$ pA | **$208.3 \pm 58.7$ pA** [109] |
| | Firing rate | $16.9 \pm 14.3$ Hz | **$16.9 \pm 14.3$ Hz** [110] |
| | Divergence | $2.8 \pm 1.9$ | **$2.8 \pm 1.9$** |
|  | Number of synapses | 406 | **406** [95] |
| | Release probability | $0.88 \pm 0.05$ | **$0.88 \pm 0.05$** [109] |
| Inhibitory (Basal) | Magnitude | $24.3 \pm 18.4$ pA | **$26.5 \pm 1.6$ pA** [109] |
| | Firing rate | $3.9 \pm 4.9$ Hz | **$3.9 \pm 4.9$ Hz** [110] |
| | Divergence | $2.7 \pm 1.6$ | **$2.7 \pm 1.6$** [111, 112] |
|  | Number of synapses | 1023 | **1023** [95] [10] |
| | Release probability | $0.72 \pm 0.10$ | **$0.72 \pm 0.10$** [109] |
| Inhibitory (Apical) | Magnitude | $24.3 \pm 33.1$ pA | **$26.5 \pm 1.6$ pA** [109] |
| | Firing rate | $3.9 \pm 4.9$ Hz | **$3.9 \pm 4.9$ Hz** [110] |
| | Divergence | $12 \pm 3$ | **$12 \pm 3$** [113][33] |
|  | Number of synapses | 1637 | **1637** [95] |
| | Release probability | $0.30 \pm 0.08$ | **$0.30 \pm 0.08$** [113] |

**Table 2.**
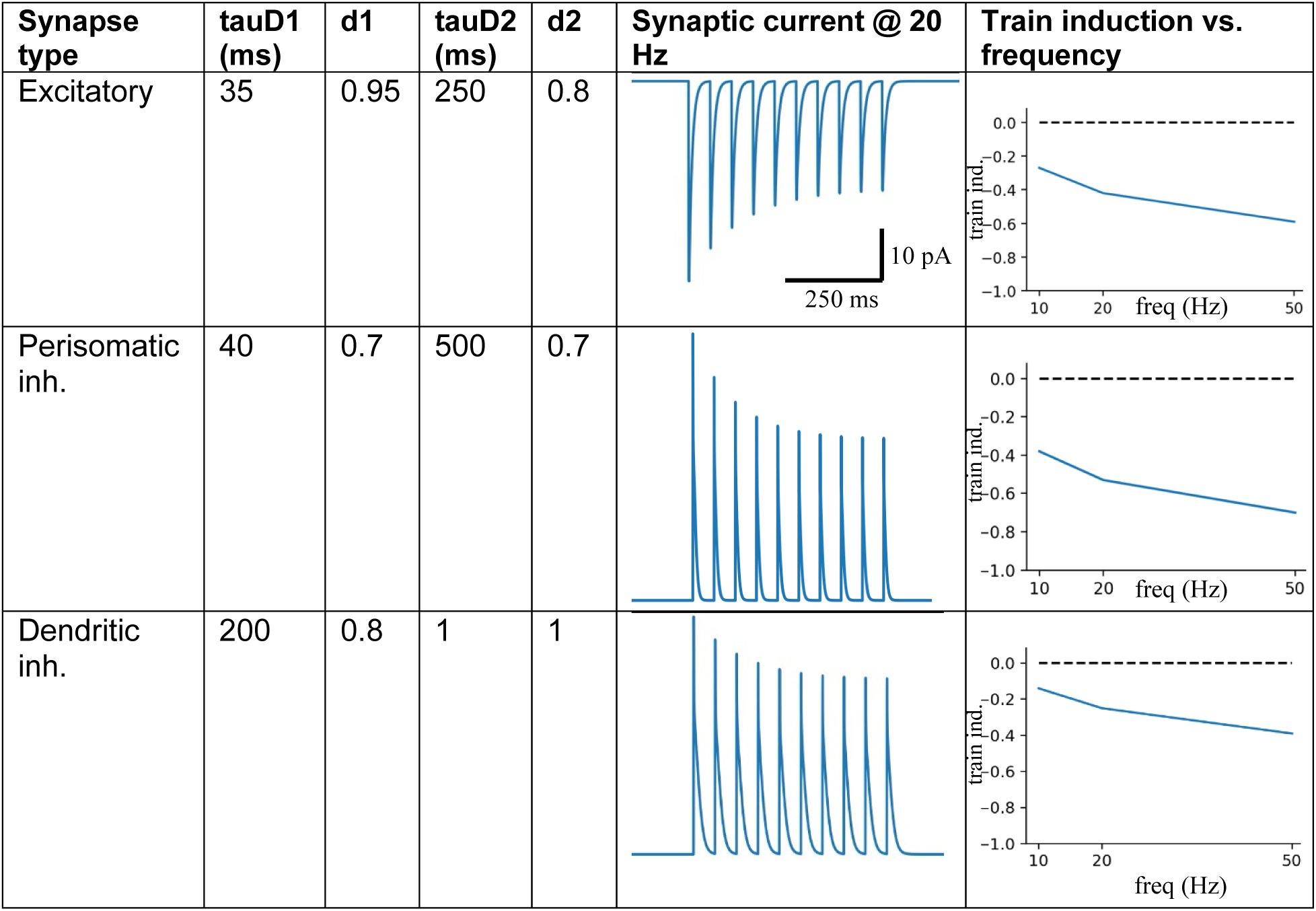
Short term pre-synaptic plasticity. Parameters were tuned to match experimental recordings reported in [76].

| Synapse type | tauD1 (ms) | d1 | tauD2 (ms) | d2 | Synaptic current @ 20 Hz | Train induction vs. frequency |
| --- | --- | --- | --- | --- | --- | --- |
| Excitatory | 35 | 0.95 | 250 | 0.8 |  |  |
| Perisomatic inh. | 40 | 0.7 | 500 | 0.7 |  |  |
| Dendritic inh. | 200 | 0.8 | 1 | 1 |  |  |

### Synapse density

Synapse density was based on a comprehensive report [95] that not only quantified the number of spines but also counted the synapses by identifying vesicle clouds and postsynaptic density using serial block-face electron microscopy. The authors found the apical tuft synapse density per micron as 2.16 ± 0.16 for excitatory and 0.22 ± 0.02 for inhibitory synapses (mean ± standard error of the mean). Other studies that only quantified spine density may undercount the true number of synapses by about 3 times because one can only count lateral spines, not those in the z-axis [96]. With that adjustment, the other values reported – 0.47 ± 0.01 spines/µm [97] and 0.53 ± 0.05 spines/µm [98]– are roughly in alignment with that in [95]. We used the values from [95] for excitatory synapse density because we found the higher value for excitatory synapse density was needed to achieve a reasonable baseline firing rate.

Morphological measurements (length of dendritic cable in µm) combined with these synapse density estimates yield the following estimates for total number of synapses: Excitatory: 16070 (2.16 syns/µm * 7440 µm) on apical dendrites and 10042 (2.16 syns/µm * 4649 µm) on basal dendrites. Inhibitory: 150 on soma [12, 99, 100] + (0.22 syns/µm *483 mm) = 256 perisomatic, 1637 (0.22 syns/µm * 7440 µm) on apical dendrites, 1023 (0.22 syns/µm * 4649 µm) on basal dendrites.

### Synaptic kinetics and short-term plasticity

All excitatory transmission was mediated by AMPA/NMDA receptors and inhibitory transmission by GABA_A_ receptors as in our prior models. The rise/decay constants for the excitatory connections were as follows: Rise/decay time constants of 0.6/6.9 ms for AMPA, and 3.7/125 ms for NMDA synapses, and a reversal potential of 0 for both. And for the inhibitory GABA-ergic connections, the rise/decay time constants were 0.5 and 6.8 ms, respectively, with a reversal potential of −75 mV. We implemented short-term depression in both the excitatory and inhibitory synapses [76] via an exponential decay of synaptic weight over short time scales [101].

Following [76] we calculated a short-term plasticity (STP) train-induction value by applying 8 pulses at 50 Hz to the synapse while in voltage clamp and measured the resulting postsynaptic current (PSC). The 6^th^, 7^th^, and 8^th^ PSCs were averaged, and the 1^st^ PSC was subtracted from the result. This value was divided by the max PSC current to yield the STP train-induction value. Excitatory synapses to L5 pyramidal neurons had an STP train-induction value of −0.34.

Inhibitory synapses arising from PV+ basket interneurons making perisomatic contact had a value of −0.60 and inhibitory synapses arising from SOM+ interneurons making dendritic contact had a value of −0.10. The equations to model the synaptic currents and short-term depression are provided in the next section.

### Modeling of synaptic currents

The corresponding synaptic currents were modeled by dual exponential functions [102, 103]. All excitatory transmission was mediated by AMPA/NMDA receptors and inhibitory transmission by GABA_A_ receptors. The corresponding synaptic currents were modeled by dual exponential functions [102, 103], as shown in eqns. 1-3,

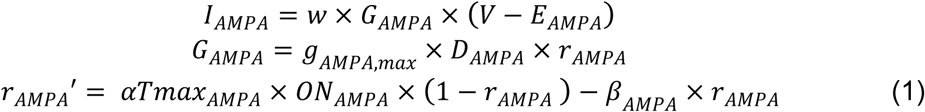

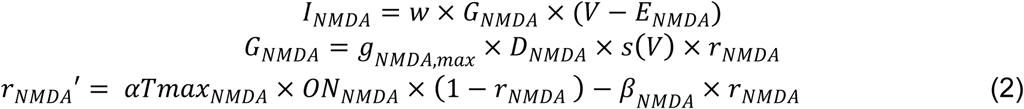

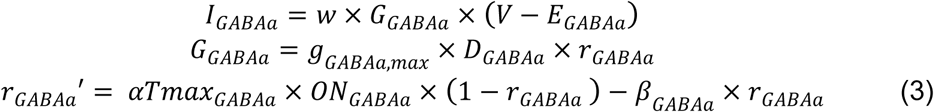

where *V* is the membrane potential (mV) of the compartment (dendrite or soma) where the synapse is located, *I* is the current injected into the compartment (nA), *G* is the synaptic conductance (µS), 𝑤𝑤 is the synaptic weight (unitless), and *E* is the reversal potential of the synapse (mV). *g_x,max_* is the maximal conductance (µS), *D* implements short-term plasticity as defined below, and *r_x_* determines the synaptic current rise and decay time constants based on the terms *αTmax* and β [102]. The voltage-dependent variable *s*(*V*) which implements the Mg^2+^ block was defined as: *s*(*V*) = [1 + 0.33 exp(−0.06 *V*)]^-1^ [104]. The terms *ON_NMDA_* and *ON_AMPA_* are set to 1 if the corresponding receptor is open, else to 0. To calculate the factor D for depression: τ_D*dD/dt=1-D and D constrained to be ≤ 1. After each stimulus, D was multiplied by a constant d (≤ 1) representing the amount of depression per presynaptic action potential and updated as D→D*d. Between stimuli, D recovered exponentially back toward 1. We modelled depression using two factors d1 and d2 with d1 being fast and d2 being slow subtypes, and d=d1*d2 was constrained to be ≥ 1.

### Synapse distribution and extrinsic inputs

We distributed 26,112 excitatory synapses over the cell. With an average of 5 synapses per pre-synaptic neuron, this yielded 5,222 presynaptic nodes. To represent pre-synaptic cell assemblies, we defined “functional groups” of presynaptic nodes. We put 100 nodes in each functional group, yielding 52 functional groups. We found that varying the number of neurons in the cell assembly did not qualitatively affect our results. To have exactly 100 nodes per functional group we rounded the total number of nodes down to 5,200. Inhibitory inputs were not defined by functional groups, but followed the literature cited in Table 1, with more than 1 synapse per pre-synaptic node. For example, with 1637 inhibitory synapses on the apical dendrites and an average of 12 synapses per pre-synaptic node, there were about 136 inhibitory nodes that contacted the apical dendrites. The numbers and distributions of model synapses along the dendrites are shown in Table 1.

The extrinsic inputs to the model were represented by spike trains arriving at the corresponding synapses along the dendritic tree. The L5 pyramidal model is spatially divided into the following sub-divisions: somatic, perisomatic, and proximal and distal components of the apical and basal trees. We used standard definitions for these sub-divisions except for perisomatic which is typically defined as the dendritic region within 100 μm of the soma [105].

<u>Tonic inputs (</u>Figs. 1-3): The firing rate of each excitatory spike train was randomly sampled from an experimentally recorded distribution (Headley, personal communication). To reproduce the 1/f spectrum that is widely found in neuronal spike trains, the 100 spike trains in each functional group were modulated by a unique normalized (from 0.5 to 1.5) firing rate time series that had a 1/f spectrum (more slow modulation of firing than fast) that was created by filtering white noise using standard techniques. The normalized firing rate time series was then multiplied with the mean firing rate of each of the 100 pre-synaptic nodes. Spike trains are then generated by sampling from a Poisson distribution using the firing rate time series of each presynaptic node of the functional group. In this way, spike trains that received the same modulation trace would be statistically correlated, representing functional presynaptic cell assemblies. Since feed-forward inhibition follows excitation in time, we averaged the firing rate time series used to design the excitatory spike trains, shifted it forward in time by 4 milliseconds, and used this to modulate the firing rate of the inhibitory spike trains.

<u>Enhanced tonic inhibition (</u>Fig. 4): Inhibitory synapses were divided into two groups, those targeting perisomatic region that originate mainly from PV+ interneurons, and those on more distal dendritic regions arising mainly from SOM+ interneurons [1, 4]. Perisomatic synapses were defined as those within 100 µm of the soma with the properties specified in Table 1 under “Inhibitory (Perisomatic and somatic).” Distal inhibitory synapses were greater than 100 µm from the soma and had their properties specified in sections “Inhibitory (Basal)” and “Inhibitory (Apical)” of the table To investigate if these inhibitory sources differentially regulated action potential initiation and dendritic spiking, we doubled the rate of presynaptic events to one of these inhibitory populations.

<u>Investigation of E/I lag (</u>Fig. 4F-I): We used the same scheme to design the extrinsic inputs as in Figs. 1-3. However, to investigate the effect of E/I lag, we increased the variance of the overall excitatory firing rate. So, while the mean rate stayed the same, we raised the modulatory trace to the fourth power so that large deviations from the mean were exaggerated. This excitatory firing rate trace was used to modulate the inhibitory firing rates (same as for Figs. 1-3). The perisomatic and distal traces were shifted independently. One was fixed at 4 ms while the other was shifted by 4, 125, 250, or 500 ms.

<u>Rhythmic inhibition (</u>Figs. 5 & 6): Gamma rhythms depend on PV+ interneurons, while beta rhythms arise from SOM+ interneurons [21]. Thus, to emulate an inhibitory rhythm we rhythmically modulated inhibitory synapses in the respective dendritic region (see description above in “Enhanced tonic inhibition”) at the appropriate frequency. Beta rhythms were emulated by modulating distal inhibitory synapses at 16 Hz with a depth of 20%. Gamma rhythms were emulated by modulating perisomatic synapses at 64 Hz with a depth of 40%. The different depths of modulation ensured that these rhythms produced similar entrainment of action potentials.

To generate inhibitory spike trains that produce rhythmic inhibition during oscillations, we modulated their collective firing rate with a sine wave:

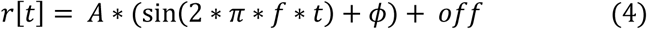

where 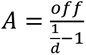 is the frequency, 𝑡𝑡 is a vector representing time, 𝜙 is the phase, 𝑜𝑓𝑓 is the offset firing rate of the spike train being modulated, and 0 < 𝑑𝑑 < 1 is the depth of modulation which represents the amplitude of the sine wave relative to 𝑜𝑜𝑓𝑓𝑓𝑓. To generate the spike train, a random vector x[t] was created with values uniformly distributed between 0 and 1. A spike was generated if *x*[𝑡] ≤ 𝑟[𝑡]𝑑𝑡 where dt in our case was 1/1000.

<u>Varying frequency of rhythmic inhibition (</u>Figs. 7 & 8): We reran the simulations used in Figures 5 and 6, but across a range of frequencies that extended above and below the beta and gamma frequencies we used in all other simulations (0.5, 2, 5, 10, 16, 20, 30, 40, 50, 60, 80 Hz).

<u>Bursty inhibition (</u>Fig. 9): We applied an envelope to the rhythmic inhibitory modulation trace to simulate the bursty temporal nature of rhythms *in vivo*. The underlying sine wave was produced in the same way as in the previous section. The envelope was a Gaussian kernel with a standard deviation of two oscillatory cycles.

<u>Clustered synaptic input (</u>Fig. 10): For this experiment we kept everything the same as in the rhythmic inhibition simulations, but introduced additional synapses driven by 40 pre-synaptic spike trains. These spike trains were created by choosing one Poisson process and jittering the spike times by ±2 ms. Each spike train drove an average of 4.6 synapses for a total of 187 synapses. These were placed either on the apical dendrites near the nexus or the basal dendrites near the soma. For both nexus and basal clustered synapses, we targeted 140 μm of dendritic cable length with 187 synapses yielding a synaptic density of 1.3 synapses per micron. For the nexus case we targeted the 140 µm region starting at the bifurcation going toward the apical dendrites. For the basal case, we targeted 3 basal dendrites that were directly connected to the soma each one had a length of 40-50 µm adding up to 140 μm total (Figure 7A). The synaptic weights were drawn from the distribution in Table 1.

### Detection of action potentials

Action potentials were detected by the NEURON software as being any time point when the somatic membrane potential crossed −10 mV with a positive first derivative.

### Detection of dendritic spikes

Na^+^ and NMDA dendritic spike events were detected following the definitions developed in [13]. We developed our own detection algorithm for Ca^2+^ spikes since they were not previously studied. Na^+^ spikes were detected if the voltage-gated sodium conductance density for a compartment exceeded 0.3 mS/cm^2^. If sodium spikes were within 2 ms of a somatic action potential, they were assumed to be a consequence of a backpropagating action potential and were not counted. An example of a Na^+^ spike can be seen in Fig. 1C1.

NMDA spikes were detected by two criteria – voltage and current. The voltage criteria were met if the membrane potential rose above −40 mV and remained there for 26 ms or more. The current criteria was met if the NMDA current exceeded 130% of the current at the −40 mV crossing and continued to be met until the current fell to 115% of that value. Whenever both the voltage and current criteria were met, an NMDA spike was recorded. An example of an NMDA spike is shown in Fig. 1C2.

Ca^2+^ spikes were detected in the same way as NMDA spikes – using a voltage and current threshold. The only difference was instead of NMDA current for the current criteria we used the sum of HVA and LVA calcium currents. An example is shown in Fig. 1C3.

### Determination of electrotonic distance

We calculated electrotonic distance using NEURON’s built-in Impedance method. The method calculates signal attenuation between two compartments at a given frequency. We used 20 Hz because the wavelength is on a time scale comparable to the membrane time constant (10s of milliseconds). However, changing this frequency 10 Hz in either direction did not qualitatively affect our results.

### Spike-triggered average (STA) of dendritic spikes (*Figs. 2*, 3)

For each dendritic compartment, we created a binary time series with 2 ms bins that was true during periods where a dendritic spike was present (based on their detected start and stop times). This was done separately for each type of dendritic spike (Na^+^, NMDA, and Ca^2+^ spikes). Each then had its spike-triggered average taken around the times of somatic action potentials and their percent change from the mean across the entire simulation calculated. The STAs were then grouped together based on their electrotonic distance percentile (10% bins) from the soma and type of dendrite (apical or basal), and the median taken for each time lag.

### Generation of F-I curves (Fig. 4)

The model with Poisson excitation and inhibition was used. In the *perisomatic* condition, we doubled the rate of inhibition to the somatic and proximal dendritic compartments (<100µm from soma). In the *distal* case, we doubled the rate of inhibition to the distal dendritic compartments (>100µm from soma). In each case, we injected 20 s current steps ranging from −1 nA to 2.4 nA in 200 pA intervals.

### Phase histogram of dendritic spikes (Fig. 5)

Entrainment of dendritic spikes to inhibitory rhythms was calculated by first converting dendritic spikes to a binary time series as was done for the STA calculation. The inhibitory rhythm was then binned by phase, with bin widths of π/4, yielding 8 phase bins. The mean of the dendritic spike binary series was taken for each phase bin. These were converted to percent change from mean and their median taken across compartments by their electronic distance and dendritic type (the same as in the STA calculation).

*Determination of action potential threshold and somatic membrane voltage distribution (Fig. 6)* Action potential threshold was taken to be the voltage 1 ms prior to the peak of the spike. Voltage distributions were calculated using 1 ms bins.

### Entrainment of dendritic spike onsets (Fig. 7)

We measured the entrainment of dendritic spike onsets to the inhibitory rhythm using the pair-wise phase consistency index, an unbiased metric [106].

### Phase histogram of dendritic spikes (Fig. 9)

For calculating entrainment of dendritic spikes to the rhythm, we followed the same procedure as detailed above for Fig. 5.

### Cross-correlation (CC) between presynaptic spikes and action potentials (Fig. 10)

We measured the effect of inhibitory rhythms on the coupling between concentrated synaptic inputs and action potentials using a phase-stratified CC. Presynaptic spike times that drove the concentrated synaptic inputs were grouped together and separated based on whether they occurred during the positive or negative phase of the imposed inhibitory rhythm. Their CC with somatic action potentials was then calculated using 1 ms bins. Since both the presynaptic and action potential times had inherent periodicities, these had to be corrected for [25]. This was performed by converting the CC and both autocorrelations to the frequency domain using the fast Fourier transform, setting the coefficients at the inhibitory modulation rhythm to 0, performing element-wise division of the CC by the square-root of the autocorrelations multiplied together, and then converting back to the time domain using the inverse fast Fourier transform on that result. To measure the strength of the cross-correlation peak, we identified the positive peak of the CC nearest to zero time lag, and then integrated over its area with the edges defined by when the CC crossed from positive to negative values.

## CODE AND DATA AVAILABILITY

Simulation code is deposited at ModelDB at https://modeldb.science/2019883. The raw simulation data are available from DBH upon request. Analysis code is posted as a GitHub repo at https://github.com/dbheadley/InhibOnDendComp.

